# Molecular characterisation of the acyltransferase-acyl carrier protein interface in a fungal highly reducing polyketide synthase

**DOI:** 10.1101/2025.10.28.685194

**Authors:** Mia E. Foran, Y. T. Candace Ho, Józef R. Lewandowski, Matthew Jenner

**Author notes:** Corresponding author: Matthew Jenner.

## Abstract

Iterative polyketide synthases (iPKSs) rely on communication between acyl carrier protein (ACP) and acyltransferase (AT) domains to ensure efficient delivery of starter and extender substrates during biosynthesis. However, the molecular determinants governing the AT:ACP interface remain poorly understood. Here, we use the fungal highly reducing PKS, SimG, a component of the cyclosporin biosynthetic pathway, as a model system to dissect the AT:ACP interface. Using alanine scanning mutagenesis combined with a high-throughput intact-protein mass spectrometry assay, we identified epitope-forming residues that affect AT:ACP interaction. These experimental constraints were used to guide docking and molecular dynamics simulations to produce a data-driven structural model of the SimG AT:ACP complex in a catalytically competent geometry. We also demonstrate that the SimG AT domain transacylates ACP domains from a range of fungal PKS architectural classes, highlighting significant interface plasticity. These insights advance our fundamental understanding of domain communication in these enigmatic megasynthases and provide a foundation for rational engineering to expand substrate scope towards novel polyketide scaffolds.

## INTRODUCTION

Fungal iterative polyketide synthases (iPKSs) are key biosynthetic enzymes responsible for producing a vast array of polyketide natural products with remarkable structural and functional diversity.^1^ These large, multidomain proteins are commonly grouped according to the extent of reductive processing that occurs during polyketide chain assembly, namely non-reducing (nrPKS), partially reducing (prPKS), and highly reducing (hrPKS) subclasses.^2,3^ Among them, hrPKSs display striking parallels to the mammalian fatty acid synthase (mFAS) in both domain organisation and catalytic strategy.^4^ Within these multifunctional enzymes, chain elongation proceeds iteratively through a series of decarboxylative Claisen condensation reactions. In each cycle, malonyl extender units, which are covalently tethered to an acyl carrier protein (ACP) domain, are used to extend the nascent polyketide chain by the ketosynthase (KS) domain. Both starter and extender units are continuously loaded onto the ACP domain by the acyl transferase (AT) domain. In contrast to mFAS, whereby the chain extended β-keto intermediate is fully reduced to a methylene by sequential activity of ketoreductase (KR), dehydratase (DH), and enoylreductase (ER) domains, hrPKSs introduce greater molecular complexity by controlling β-carbon processing. This allows for the incorporation of β-hydroxyl groups (via KR alone), α,β-unsaturated double bonds (via KR and DH), or saturated methylene units (via KR, DH, and ER). Additionally, some hrPKSs include a functional methyltransferase (MT) domain capable of α-carbon methylation, an enzymatic step known to influence biosynthetic programming, but is not active in mFAS. Throughout this process, intermediates and extender units remain covalently attached via a thioester bond to a 4’-phosphopantetheine (Ppant) prosthetic group, which is post-translationally attached to the ACP domain (**Fig. 1a**).

**Figure 1.**
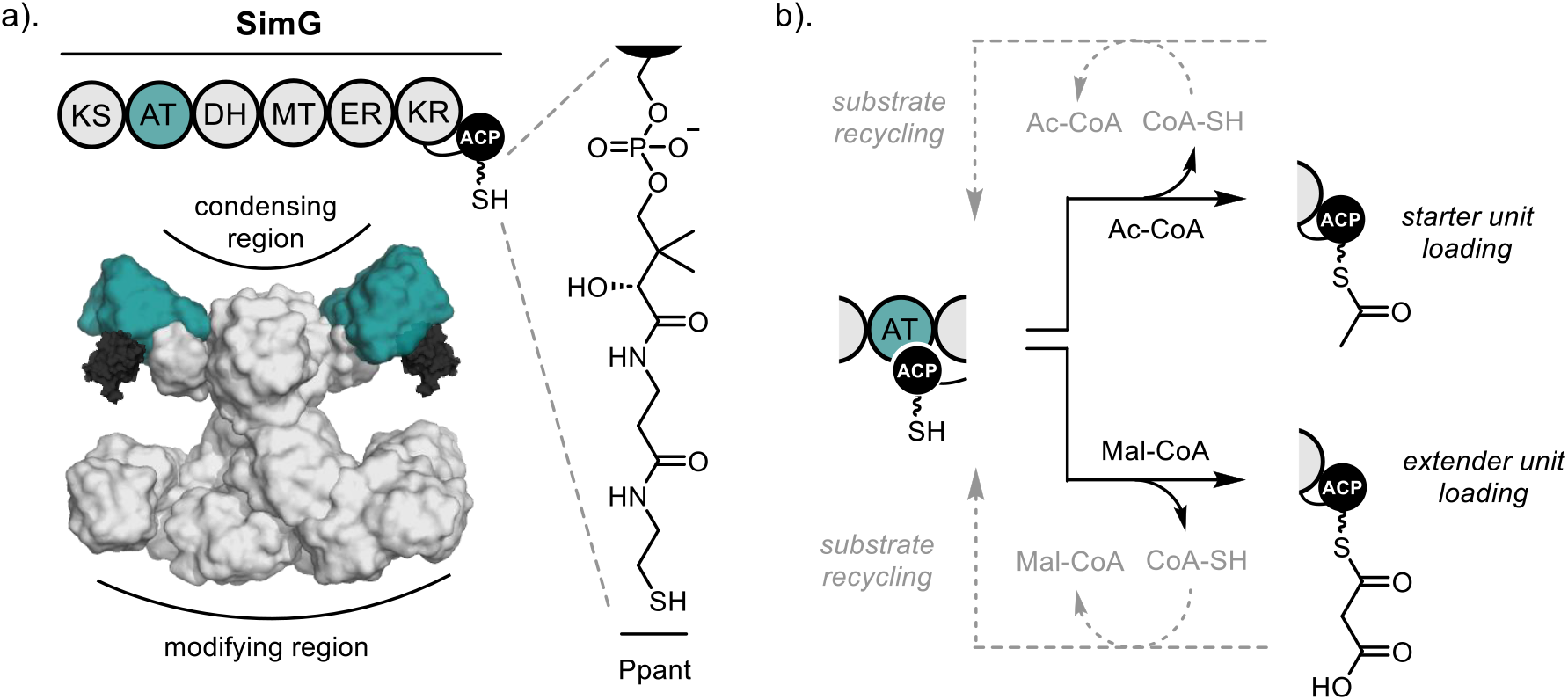
Role of the acyltransferase domain in fungal hrPKSs. a). Domain organisation (*top*) and structural model (*bottom*) of the SimG hrPKS responsible for production of *E*-2-butenyl-4-methyl-threonine (Bmt) incorporated into cyclosporin A. The AT and ACP domains are highlighted in teal and black, respectively, and the structure of the Ppant group is shown. b). Schematic overview of AT catalysis, including acetyl /malonyl transacylation onto the ACP domain (black arrows) and substrate recycling back to CoA (grey dashed arrows).

In fungal hrPKSs, the AT domain plays a pivotal role in the biosynthetic process, exhibiting dual specificity to load both starter and extender units.^5,6^ While acetyl-CoA is the most common starter unit, exceptions include propionyl-CoA, benzoyl-CoA, and nicotinyl-CoA,^7–10^ with malonyl-CoA serving exclusively as the extender unit. During biosynthesis, the AT domain catalyses a two-step transfer of a given acyl group from CoA to the Ppant arm of the ACP domain. Initially, an active site Ser residue is acylated by the acyl-CoA, followed by transfer of the acyl group to the Ppant thiol of the ACP domain, with both steps being reversible. Due to its dual specificity, the AT domain must discriminate between acetyl- and malonyl-CoA in the cellular environment, where these substrates can act as competitive inhibitors of one another. Our previous work has shown that fungal hrPKSs employ a substrate recycling mechanism, similar to mFAS, to overcome this issue. Here, the inherent reversibility of the reaction is harnessed, allowing incorrectly loaded acyl groups to be returned back to CoA, ensuring biosynthetic fidelity of the system (**Fig. 1b**).^6,11^

Despite the fundamental role of the AT domain, molecular details underpinning the AT:ACP interaction in fungal hrPKSs remain poorly understood. A key challenge is their dynamic and transient nature, which makes them difficult to capture using conventional structural techniques. Although the natural tethering of domains increases the effective local concentration of ACP^12^, the resulting interactions are typically low affinity and thus hard to characterise experimentally. Recent cryo-EM structures of the LovB hrPKS were unable to resolve the ACP domain^13^, while related work on mFAS has captured density for the ACP in various states as it shuttles between KS and AT domains.^14^ These structural snapshots underscore the complex choreography of ACP domain movement required for processivity. However, notwithstanding these insights, critical aspects such as the precise interface with the AT domain and the conformation of the Ppant arm remain unresolved, leaving a significant gap in our understanding of AT:ACP recognition. This information has significant implications for engineering efforts. In particular, if individual AT domains, or indeed entire condensing regions, are to be swapped between systems to create novel biosynthetic pathways, it is essential to determine whether native AT:ACP interactions can be preserved or re-established. A detailed understanding of the molecular determinants that mediate this interaction is therefore a key step toward rational reprogramming of polyketide biosynthesis.

In this study, we focus on the SimG hrPKS as a model system to dissect the AT:ACP interface. Using a alanine scanning mutagenesis approach, in conjunction with a high-throughput intact protein mass spectrometry assay, we have identified critical residues involved in mediating the AT:ACP domain interaction. These experimental data were used to guide protein docking and molecular dynamics simulations to generate a model of the SimG ACP:AT complex in a catalytically competent state. Importantly, we also show that the interface exhibits functional plasticity, as the AT domain is capable of transacylating a range of ACP domains across different hrPKS classes. Together, our findings provide a foundation for understanding the specificity and flexibility of ACP:AT interactions, with direct implications for the future design of engineered hrPKSs.

## RESULTS & DISCUSSION

### Mapping the AT-binding epitope for SimG ACP domain

Our previous work on the SimG condensing region established an intact protein mass spectrometry-based assay to monitor AT-catalysed malonyl transfer from CoA to the *holo*-ACP domain.^6^ A time-resolved analysis of this assay showed that the reaction reaches near-completion, whilst still in the linear region, after 2 minutes (**Supplementary Fig. S1**). This time-point was therefore used for all subsequent experiments to observe changes in reaction efficiency. In order to map the AT-binding epitope of the SimG ACP domain, we employed alanine scanning mutagenesis; a proven method for identifying surface interaction sites on carrier proteins.^15–19^ Using an AlphaFold model of the SimG ACP domain, we identified 47 surface-exposed residues, which were individually mutated to alanine (X→Ala) to eliminate side-chain functionality (**Supplementary Fig. S2**). All mutants were subsequently overproduced and purified to homogeneity in their *apo*-ACP form. Of these, five mutants (K2477A, K2478A, D2513A, H2545A, and E2559A) gave poor protein yields, while four others (F2501A, L2503A, S2542A, and M2544A) were entirely insoluble. In addition, mass spectrometry analyses revealed apparent degradation in six additional soluble mutants (E2475A, I2481A, K2484A, D2504A, P2506A, and S2558A). This process resulted in a final set of 32 soluble, stable X→Ala surface mutants, collectively representing ∼ 49 % of the solvent-exposed (non-Gly/Ala) surface residues of the SimG ACP domain from the core helical region. Mutants were enzymatically converted to their *holo*-form and subjected to intact protein mass spectrometry, which confirmed efficient phosphopantetheinylation.

The library of *holo*-SimG ACP X→Ala mutants was subjected to our mass spectrometry-based malonyl transfer assay to evaluate the contribution of individual residues to the AT:ACP interaction interface. Here, malonyl transfer efficiency was quantified by calculating the ratio of malonyl-ACP to *holo*-ACP signal and expressing this value as a percentage relative to wild-type SimG ACP activity. While most X→Ala mutants did not replicate wild-type activity precisely, the majority retained 60 – 90 % of wild-type malonyl transfer, suggesting that alanine substitution at these positions does not significantly disrupt interaction with the SimG AT domain. In contrast, six mutants (L2494A, I2495A, K2496A, K2505A, N2522A, and E2531A) exhibited markedly reduced activity (≤ 50 %), consistent with a more significant contribution for these residues in AT binding (**Fig. 2a, Supplementary Fig. S3**). Circular dichroism spectroscopy of the four most disruptive mutants revealed spectra nearly identical to that of wild-type SimG ACP, indicating that the observed reduction in activity is unlikely due to altered secondary structure or unfolding (**Supplementary Fig. S4**). Instead, the loss of function likely stems from disruption of critical side chain interactions at the AT:ACP interface. These results implicate the six identified residues as contributors to AT:ACP binding, albeit to varying degrees. Mapping these positions onto an AlphaFold model of SimG ACP domain revealed that four key residues (L2494, I2495, K2496, and K2505) are clustered within the loop connecting helix I to helix II’. The remaining two, N2522 and E2531, are located on helix II and the loop between helix II and III, respectively (**Fig. 2b**, *top*). This spatial distribution defines a focused interaction epitope on the ACP domain surface proximal to the Ppant attachment site (**Fig. 2b**, *bottom*), indicating that these residues are situated in a functionally significant region.

**Figure 2.**
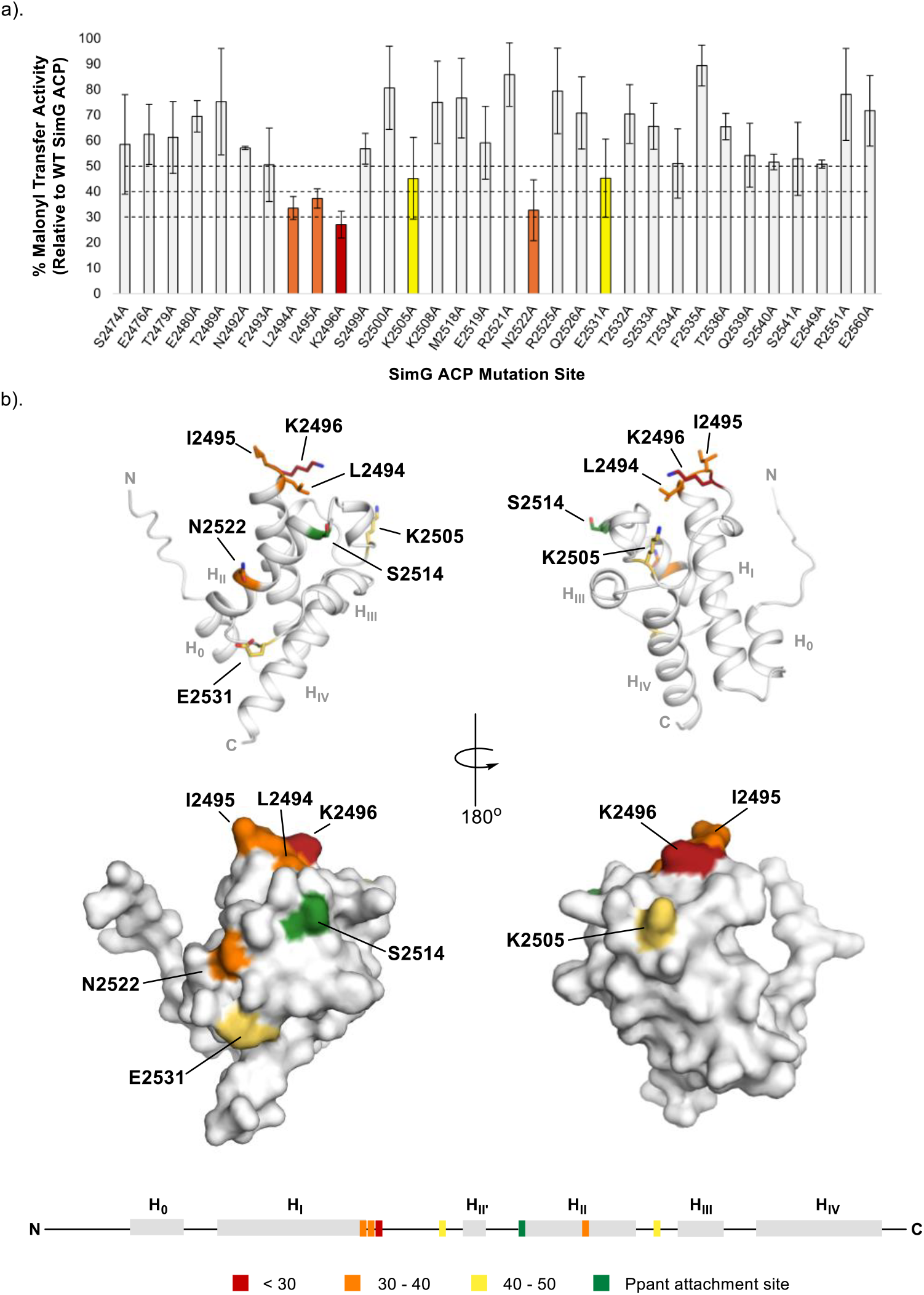
Mapping the AT-binding epitope on the SimG ACP domain. a). Bar chart showing SimG AT-catalysed malonyl transfer activity for all X → Ala of the SimG ACP domain. Data is plotted as a percentage of wild-type SimG ACP activity after a 2 min incubation period. Arbitrary cut-off bounds for reduction in transacylation activity are applied and highlighted in grey (> 50 %), yellow (40 – 50 %), orange (30 – 40 %), and red (< 30 %). Error bars represent ± 1 standard deviation (1σ) from the mean, where *n* = 3. b). Relative transacylation activities of X → Ala mutants mapped onto an AlphaFold model of the SimG ACP domain, and colour-coded according to the accompanying bar chart. Cartoon (*top*) and surface (*bottom*) representations are shown. Residues with activities in the 40 – 50 %, 30 – 40 %, and < 30 % ranges are highlighted in stick representation. A linear schematic of the SimG ACP domain secondary structure is included, indicating residues critical for interaction with the SimG AT domain. The site of Ppant attachment (Ser2514) is highlighted in green.

### Elucidating the ACP-binding epitope on SimG AT domain

Having identified the key ACP residues mediating interaction with the AT domain, we next focused on the AT domain itself. Due to its size (51 kDa), a comprehensive surface X→Ala scan of the AT domain was not feasible. Instead, a targeted mutagenesis approach was employed, focusing on 10 surface-exposed residues surrounding to the Ppant-binding region and catalytic chamber (**Supplementary Fig. S5**). Prior to evaluating ACP transacylation, each AT mutant was assayed for self-acylation at the catalytic Ser706 using malonyl-CoA. Given the proximity of the selected residues to the CoA /Ppant-binding channel and active site, mutations at these positions have the potential to disrupt substrate binding and acylation, which could confound interpretation of transacylation results by mimicking effects attributable to an altered ACP interaction. Acylation of the AT domain was conducted using a 5-fold excess of malonyl-CoA, with the majority of mutants exhibiting acylation levels comparable to wild-type. However, the S771A variant displayed < 50 % of wild-type activity, implicating this residue in CoA recognition (**Supplementary Fig. S6a**). Structural overlay of the SimG AT AlphaFold model with the crystal structure of the *E. coli* FAS AT:malonyl-CoA complex (PDB: 2G2Z) allowed an approximation of CoA-binding in the SimG AT cavity. Here, spatial proximity between Ser771 and the NH_2_ group of the CoA adenosine moiety allow for the possibility of a hydrogen bonding interaction (**Supplementary Fig. S6b**). Due to the significantly reduced ability to be transacylated, this mutant was therefore excluded from subsequent analyses.

The remaining nine AT variants were evaluated for ACP transacylation, from which three mutants (R798A, R801A, and R915A) retained wild-type activity. Moderate reductions (60 – 80 %) were observed for F796A and E892A, while Q769A and N771A exhibited ≤ 30% activity, and Q897A and R900A showed ≤ 20 % (**Fig. 3a**). Mapping these activity profiles onto the SimG AT AlphaFold model highlighted that residues which perturbed ACP recognition (Q770, N772, E893, Q898 and R901) are located at the top of the substrate binding cleft, whilst those lining the flanks (R799, R802 and R916) had minimal effect (**Fig. 3b**). This location is congruent with positioning of the carrier protein in structures of covalently trapped AT:ACP complexes from different systems.^20–22^

**Figure 3.**
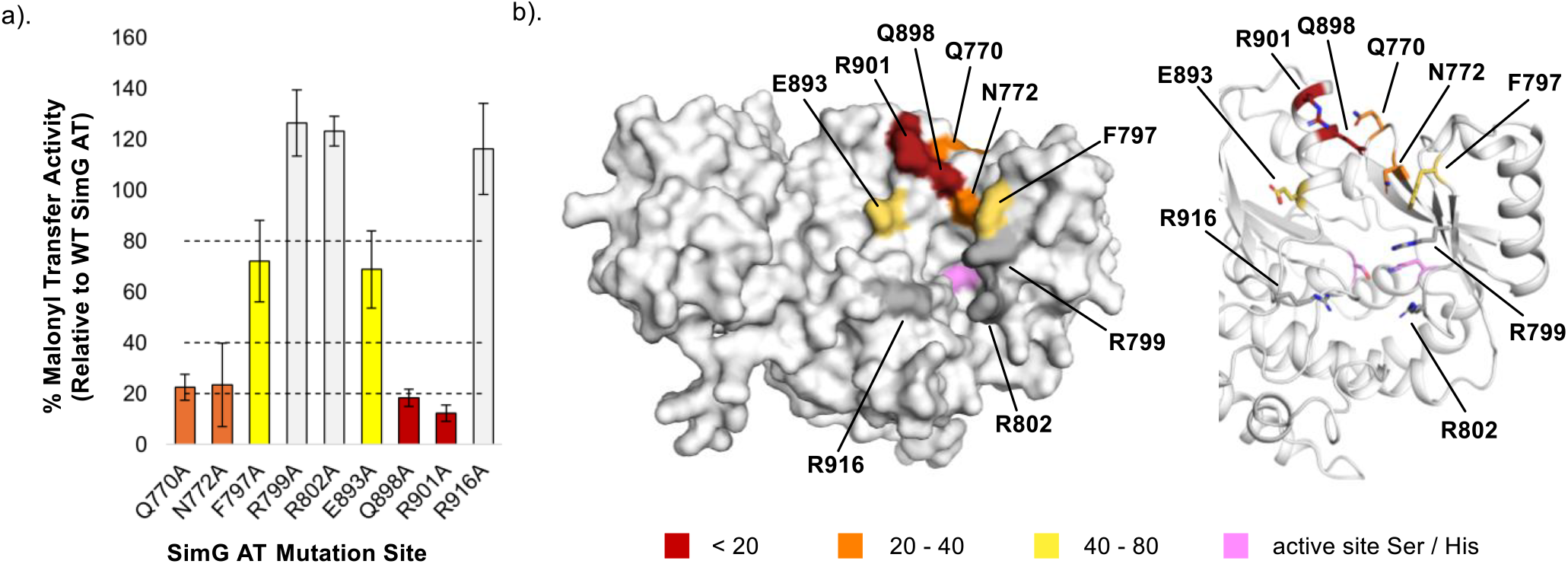
Mapping the ACP-binding epitope on the SimG AT domain. a). Bar chart showing relative malonyl transfer activity for all X → Ala mutants of the SimG AT domain. Data is plotted as a percentage of wild-type SimG AT activity after a 2 min incubation period. Arbitrary cut-off bounds for reduction in transacylation activity are applied and highlighted in grey (> 80 %), yellow (40 – 80 %), orange (20 – 40 %), and red (< 20 %). Error bars represent ± 1 standard deviation (1σ) from the mean, where *n* = 3. b). AlphaFold model of the SimG AT domain, with relative transacylation activities of X → Ala mutants mapped onto an and colour-coded according to the bar chart. Surface (*left*) and cartoon (*right*) representations are shown, and the catalytic Ser706 and His808 residues are highlighted in pink.

### Computational docking and validation of the SimG AT:ACP complex

With residue-level information of the SimG AT:ACP epitope in hand, these experimentally-derived insights were used as restraints for HADDOCK docking simulations (keeping Ser2514 as a passive residue)^23^, enabling the construction of a model guided by our observations. Using AlphaFold-predicted structures of the individual AT and ACP domains as inputs^24^, the resulting model of the SimG AT:ACP complex identified a top-scoring solution that was highly consistent with the experimental data and with high complimentary for shape and charge. In this model, Ser2514 was positioned at the entrance of the substrate-binding cleft, approximately 15 - 16 Å from the catalytic Ser706 residue of the AT domain. It is worth highlighting that attempts to model the AT:ACP domain complex with AlphaFold produced non-sensical solutions, with the ACP domain docked on the opposite face to the substrate binding cleft (**Supplementary Fig. S7**).

Using this initial model, a malonyl moiety was then manually appended to Ser706 of the AT domain, guided by preexisting structures of malonyl-bound AT domains^25,26^, and a Ppant arm was covalently tethered to Ser2514 of the ACP domain and manually modelled into the structure in an arbitrary extended conformation. The assembled complex was subjected to energy minimisation followed by classical molecular dynamics (cMD) simulations for 250 ns to capture potential conformational changes. Throughout the simulations, geometric parameters relevant to malonyl group transfer were monitored, including the distance between the Ppant thiol and the histidine nitrogen (His808[**Nε**] → Ppant—[S—**H**]), and the distance and angle between the Ppant thiol and the carbonyl carbon of the malonyl group (Ppant—[**S**—H] → Mal[**C1**]—Ser706), which together reflect catalytic readiness. Throughout the simulation, numerous frames met one or more of the parameters (His808[**Nε**] → Ppant—[S—**H**] distance <2.5 Å; Ppant—[**S**—H] → Mal[**C1**] → Mal[**O**]—Ser706 angle: 100 – 110° i.e., approaching the Bürgi-Dunitz angle^27^, Ppant—[**S**—H] → Mal[**C1**]—Ser706 distance < 4 Å) required for a catalytically competent state, suggesting that the docked complex is valid. However, no single frame met all three criteria perfectly, suggesting that transfer of the malonyl unit may not be concerted, rather a tightly coupled process requiring some small movements in the active site. In general, for frames where the conditions for thiol proton abstraction by His808 are more favourable, the geometry is less optimal for the nucleophilic attack of the sulphur on the carbonyl, and vice versa.

In a representative frame (**Fig. 4a**), a plausible geometry for both abstraction of the thiol proton and nucleophilic attack are achieved with the exception that the malonyl carbonyl is not fully seated in the oxyanion hole formed by the backbone amide of Ser707 and Gln618. Here, the carbonyl is hydrogen bonded to the amide proton of Ser707, but 3.8 Å away from amide proton of Gln618. In the presented frame, the ACP binds to the AT domain in essentially the same orientation as in the initial docked model. In this conformation, K2496 of ACP domain, whose X→Ala mutant has the strongest effect on malonyl transfer efficiency, forms electrostatic interactions with E893 of the AT domain (**Fig. 4b**). Additionally, I2495 of ACP domain packs against a hydrophobic patch formed by L900, Y910, L883 and V896 (**Fig. 4c**), which is locked into position by L2494. Interestingly, N2522 and K2505 are not engaged in any stabilising interactions in the presented frame, despite having a notable effect on malonyl transfer. During the simulations, whilst the ACP domain maintains a generally consistent orientation and position relative to the AT domain, it appears to sample the interface near the entrance of the AT channel leading to the active site, rather than remaining fixed in a single set of interactions, similar to observations made for the *E. coli* FabD:AcpP complex.^22^ To assess whether residues N2522 and K2505 participate in transient contacts, we extended the classical MD simulation with an additional 200 ns of accelerated MD. During the simulation, N2522 frequently forms hydrogen bonds with G795, located in a loop region of the AT domain. Likewise, K2505 exhibits transient interactions with D595 on the opposite side of the interface. Comparison of our complex with a recent cryo-EM structure of mFAS, in which partial density for the ACP was observed at the AT domain, indicates that the positioning of the ACP domain in our docked complex is consistent with that observed experimentally (**Supplementary Fig. S8**).^14^ The lack of well-defined density for ACP in cryo-EM map is also consistent with the ACP sampling the interface with MAT as observed in our MD simulations.

**Figure 4.**
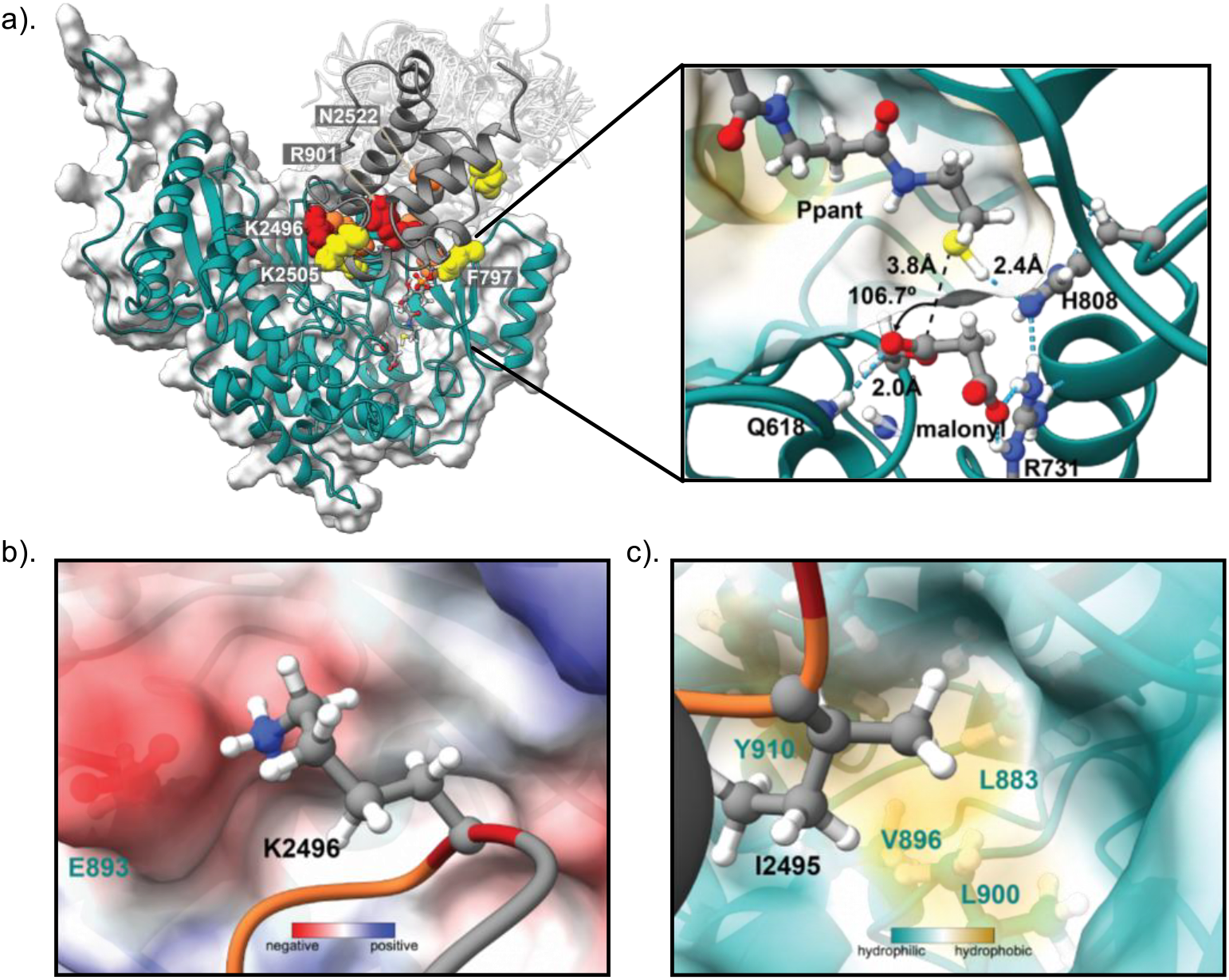
Docked complex of malonyl-SimG AT domain with *holo*-SimG ACP domain. a). Representative frame from a classical MD simulation starting from a docked ACP:AT complex, where Ppant is primed for reaction with malonyl. Surface residues identified by alanine scanning mutagenesis as being important for productive binding are shown as spheres with colours indicating malonyl transfer efficiency as in **Fig. 2** and **3**. White semi-transparent cartoons depict the range of conformations sampled by the complex in a 200 ns aMD simulation. The SimG AT and ACP domains are displayed in teal and grey cartoon, respectively. A magnified view of the substrate binding pocket from the MD frame is shown, with key distances (black dashed lines) and the Ppant—[S—H] → Mal[C1] → Mal[O]—Ser706 angle highlighted. Cyan dashed lines indicate selected hydrogen bonds. b). Electrostatic interaction between K2496 of the ACP domain and E893 of the AT domain. c). Insertion of I2495 from the ACP domain into a hydrophobic cleft on the AT domain surface formed by residues L883, V896, L900, and Y910. The co-ordinates for the frame shown in panel (a), along with the cMD /aMD trajectories stripped of water and ions with frames every 20 ns are deposited at https://doi.org/10.5281/zenodo.17272938.

### Probing carrier protein tolerance of the SimG AT domain

Having established a model for the SimG AT:ACP complex, we next examined the capacity of the AT domain to functionally interact with noncognate carrier protein domains. Here, we selected a range of carrier proteins to reflect the various commonly occurring hrPKS domain architectures. These include the canonical hrPKS domain arrangement with an active integrated ER domain, termed *cis*-ER (e.g. LovF, lovastatin biosynthesis^28,29^), or with an inactive integrated ER domain, which is supplemented by a stand-alone ER domain, termed *trans*-ER (e.g LovB, lovastatin biosynthesis^28,30^). In addition, we examined carrier proteins from a dual hrPKS-nrPKS system (e.g. CazF-CazM, chaetoviridin /chaetomugilin biosynthesis^31^) and from a hybrid hrPKS-nonribosomal peptide synthetase (NRPS) (e.g. CcsA, cytochalasin biosynthesis^32^). In the latter, both the ACP and peptidyl carrier protein (PCP) domains were considered due to the covalent tethering of the system (**Fig. 5a**).

**Figure 5.**
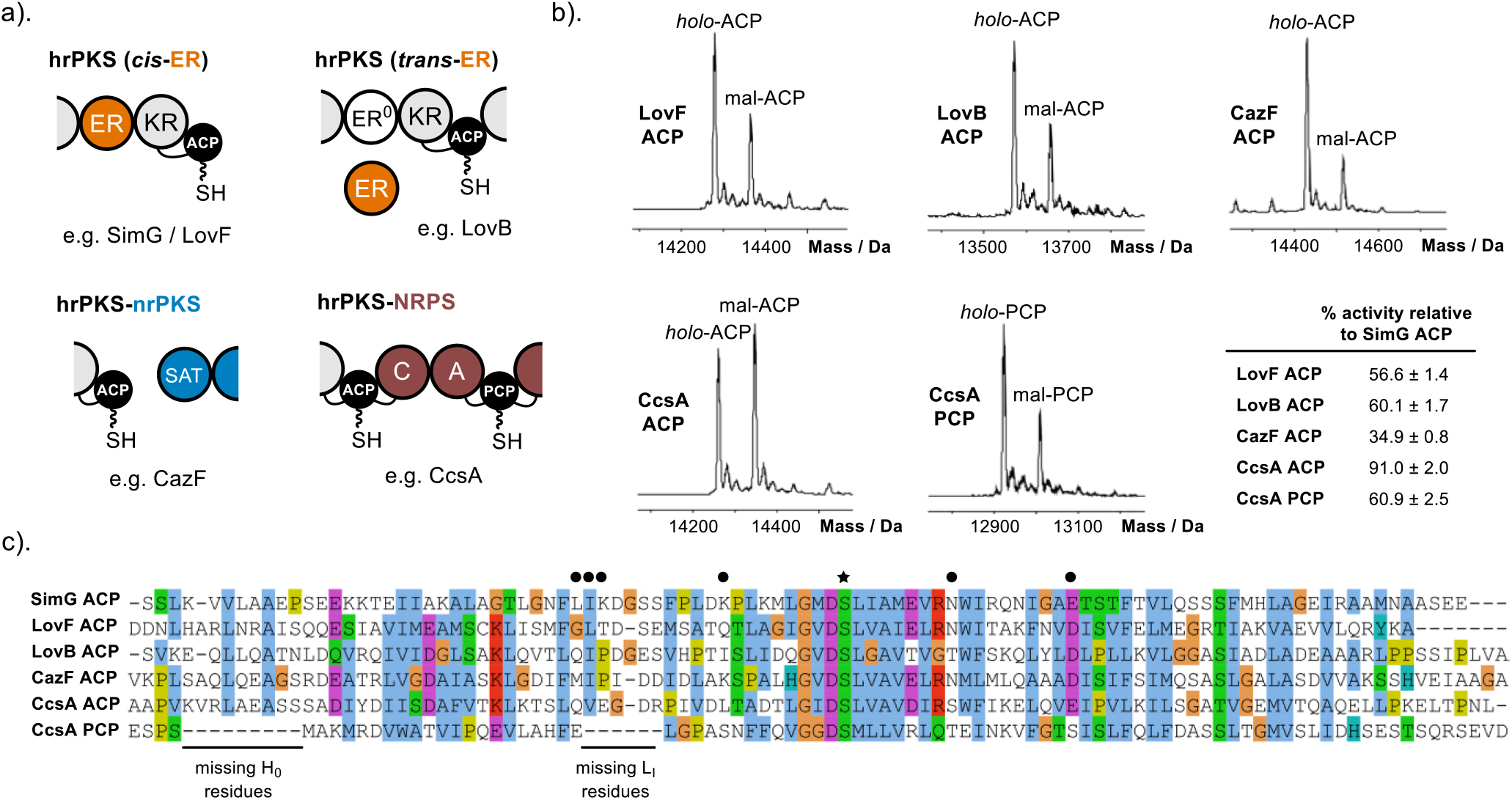
Investigating carrier protein specificity of the SimG AT domain. a). Domain architectures of different hrPKS classes. The defining features in each case are highlighted – orange, *cis*-/*trans*-ER; blue, SAT domain of nrPKS machinery; red, catalytic domains and PCP domain of NRPS machinery. b). Representative deconvoluted mass spectra of LovF ACP, LovB ACP, CazF ACP, CcsA ACP and CcsA PCP domains following incubation with SimG AT and malonyl-CoA for 2 min. The peaks for *holo*- and malonyl-carrier protein forms are labelled and the amount of transacylation relative to the cognate SimG ACP is stated (see **Supplementary Fig. S3**, *top*). The errors reported represent ± 1 standard deviation (1σ) from the mean, where *n* = 3. c). Multiple sequence alignment of all carrier proteins tested for transacylation with SimG AT domain. The missing residues in the L_I_ region of CcsA PCP domain are highlighted. Residues identified as important for AT-binding in the SimG AT:ACP complex are annotated with black circles, and the Ppant attachment site (Ser2514) with a black star.

Carrier protein domains excised from each system were tested in our malonyl transfer assay to evaluate cross-compatibility with the SimG AT domain (**Supplementary Fig. S9**). Under the same assay conditions outlined previously, all carrier protein domains were loaded with malonyl to varying degrees, although none to the same level as SimG ACP, highlighting notable interfacial plasticity whilst still displaying a preference for the cognate ACP domain (**Fig. 5b**). At the sequence level it is evident that key residues identified in the L_I_ region have a degree of hydrophobic conservation (F/L-X-L/I/V) (**Fig. 5c**), likely allowing these ACP domains to interact productively with the hydrophobic pocket identified during MD simulations (**Fig. 4c**). Among the ACP domains tested, CcsA ACP exhibited activity levels comparable to those of the cognate SimG ACP. Sequence and structural analyses show that CcsA ACP possesses an Arg residue in the Loop I region, positioned four residues downstream from the critical K2496 in SimG ACP (**Fig. 5c**). This Arg residue likely mimics the positive charge in Loop I required for efficient malonyl transfer, enabling interaction with a cluster of acidic residues on the AT domain surface that form a negatively charged ridge (**Supplementary Fig. S10a**). Notably, the other ACP domains tested lack a positively charged residue in the early portion of Loop I and instead contain several acidic residues in this region (**Supplementary Fig. 10b**), contributing to their reduced malonyl transfer activity.

The CazF ACP domain exhibited the lowest levels of acylation, and harbours three Asp residues in the Loop I region, likely disfavouring interaction with the acidic patch on SimG AT (**Supplementary Fig. 10b**). However, this feature in combination with other specific structural characteristics, may be critical for it to engage in inter-subunit communication with a starter unit AT (SAT) domain from an nrPKS system.^33^ Interestingly, the carrier proteins from CcsA showed different levels of acylation, with the ACP domain preferentially loaded over the PCP domain. Sequence analysis reveals the PCP domain lacks six L_I_ region residues present in ACP domains and implicated in AT binding (**Fig. 5c**). Structurally, this deletion shortens the L_I_ segment in the PCP domain relative to its ACP counterpart and appears to also form a short helix in this region when modelled with AlphaFold (**Supplementary Fig. S11a**). The different levels of malonyl transfer observed might be a requirement of the covalently tethered hrPKS-NRPS hybrid system, reducing PCP acylation by the AT domain and ensuring biosynthetic fidelity. Indeed, AlphaFold modelling of CcsA suggests that flexible linkers could bring the PCP domain into proximity with the AT domain (**Supplementary Fig. S11b**), where loading with acetyl /malonyl would obstruct L-Phe incorporation by the NRPS, effectively halting the assembly line.

## CONCLUSIONS

Communication between the AT and ACP domains is fundamental to the function of fungal hrPKSs, ensuring efficient loading of starter and extender units during biosynthesis. Here, we provide a residue-level characterisation of the AT:ACP interface from the SimG hrPKS, establishing a framework for understanding domain communication within these iterative megasynthases. Our data highlights specific electrostatic and hydrophobic interactions that contribute to complex formation, while also revealing inherent plasticity at the interface. No single point mutation was sufficient to abolish activity, and our MD simulations suggest that the ACP domain can adopt multiple productive binding modes at the AT domain surface. These features likely help the interface tolerate naturally occurring mutations, while also enabling the successful evolutionary diversification of hrPKSs over time.

Notably, the SimG AT domain was able to productively interact with ACP domains from non-cognate hrPKS systems, indicating that key elements of the interface are maintained across hrPKSs. Although all ACP domains possessed residues that could contribute to the hydrophobic interface region, improvements in malonyl transfer could be mapped to the presence of a basic residue that compliments an acidic patch on the AT domain. It is interesting to consider that the remaining sequence variations between ACP domains (see **Fig. 5c**) may therefore reflect functional specialisation, potentially linked to interactions with stand-alone enzymes, partner subunits, or indeed specific programming requirements. The observed flexibility in ACP domain recognition undoubtedly opens promising opportunities for rational engineering of hrPKSs; this could include the exchange of AT domains or entire condensing regions between systems, with confidence that productive AT:ACP communication can be retained.

## ACKNOWLEDGEMENTS

This work was supported by a UKRI Future Leaders Fellowship to M.J. (MR/W011247/1), from which Y.T.C.H was funded. The Bruker MaXis II instrument used in this study was funded by the BBSRC (BB/M017982/1). M.E.F gratefully acknowledges funding from an EPSRC Doctoral Training Partnership (EP/T51794X/1) studentship. J.R.L. acknowledges funding from BBSRC: BB/Z517318/1, UKRI723 and UKRI718, EPSRC: EP/Z531200/1 and EP/Z535709/1, and MRC UKRI877. The computing facilities were provided by the Scientific Computing Research Technology Platform of the University of Warwick. The authors are grateful to Professor Yifan Chen and Dr. Wooyoung Choi for providing their docked model of the mFAS AT:ACP complex for comparative analysis, and Dr. Nikola Chmel for assistance with circular dichroism measurements.

## AUTHOR CONTRIBUTIONS

M.J. and M.E.F. conceived and designed the study. M.E.F. constructed all SimG ACP and AT mutant plasmids. M.E.F and Y.T.C.H. carried out protein overexpression and purifications, and biochemical assays on recombinant proteins. M.E.F. J.R.L. and M.J. generated AlphaFold models, and J.R.L. conducted docking studies and molecular dynamics simulations of the protein interface. M.J. wrote the manuscript with input from all authors.

## REFERENCES

(1) Chooi, Y. H.; Tang, Y. Navigating the Fungal Polyketide Chemical Space: From Genes to Molecules. J. Org. Chem., 2012, 77, 9933–9953.

(2) Cox, R. J. Polyketides, Proteins and Genes in Fungi: Programmed Nano-Machines Begin to Reveal Their Secrets. Org. Biomol. Chem. 2007, 5, 2010–2026.

(3) Cox, R. J. Curiouser and Curiouser: Progress in Understanding the Programming of Iterative Highly-Reducing Polyketide Synthases. Nat. Prod. Rep. 2023, 40, 9–27.

(4) Herbst, D. A.; Townsend, C. A.; Maier, T. The Architectures of Iterative Type I PKS and FAS. Natural Product Reports, 2018, 35, 1046–1069.

(5) Ma, S. M.; Tang, Y. Biochemical Characterization of the Minimal Polyketide Synthase Domains in the Lovastatin Nonaketide Synthase LovB. FEBS J. 2007, 274, 2854–2864.

(6) Foran, M. E.; Auckloo, N. B.; Ho, Y. T. C.; Liu, S.; Hai, Y.; Jenner, M. Biochemical Dissection of a Fungal Highly Reducing Polyketide Synthase Condensing Region Reveals Basis for Acyl Group Selection. Chem. Sci. 2025, 16, 13173–13182.

(7) Zou, Y.; Xu, W.; Tsunematsu, Y.; Tang, M.; Watanabe, K.; Tang, Y. Methylation-Dependent Acyl Transfer between Polyketide Synthase and Nonribosomal Peptide Synthetase Modules in Fungal Natural Product Biosynthesis. Org. Lett. 2014, 16, 6390–6393.

(8) Iqbal, Z.; Han, L.-C.; Soares-Sello, A. M.; Nofiani, R.; Thormann, G.; Zeeck, A.; Cox, R. J.; Willis, C. L.; Simpson, T. J. Investigations into the Biosynthesis of the Antifungal Strobilurins. Org. Biomol. Chem. 2018, 16, 5524–5532.

(9) Bonsch, B.; Belt, V.; Bartel, C.; Duensing, N.; Koziol, M.; Lazarus, C. M.; Bailey, A. M.; Simpson, T. J.; Cox, R. J. Identification of Genes Encoding Squalestatin S1 Biosynthesis and in Vitro Production of New Squalestatin Analogues. Chem. Commun. 2016, 52, 6777–6780.

(10) Itoh, T.; Tokunaga, K.; Matsuda, Y.; Fujii, I.; Abe, I.; Ebizuka, Y.; Kushiro, T. Reconstitution of a Fungal Meroterpenoid Biosynthesis Reveals the Involvement of a Novel Family of Terpene Cyclases. Nat. Chem. 2010, 2, 858–864.

(11) Stern, A.; Sedgwick, B.; Smith, S. The Free Coenzyme A Requirement of Animal Fatty Acid Synthetase. Participation in the Continuous Exchange of Acetyl and Malonyl Moieties between Coenzyme A Thioester and Enzyme. J. Biol. Chem. 1982, 257, 799–803.

(12) Kuriyan, J.; Eisenberg, D. The Origin of Protein Interactions and Allostery in Colocalization. Nature, 2007, 450, 983–990.

(13) Wang, J.; Liang, J.; Chen, L.; Zhang, W.; Kong, L.; Peng, C.; Su, C.; Tang, Y.; Deng, Z.; Wang, Z. Structural Basis for the Biosynthesis of Lovastatin. Nat. Commun. 2021, 12, 867.

(14) Choi, W.; Li, C.; Chen, Y.; Wang, Y.; Cheng, Y. Structural Dynamics of Human Fatty Acid Synthase in the Condensing Cycle. Nature 2025, 641, 529–536.

(15) Wong, F. T.; Jin, X.; Mathews, I. I.; Cane, D. E.; Khosla, C. Structure and Mechanism of the trans-Acting Acyltransferase from the Disorazole Synthase. Biochemistry, 2011, 50, 6539–6548.

(16) Ye, Z.; Musiol, E. M.; Weber, T.; Williams, G. J. Reprogramming Acyl Carrier Protein Interactions of an Acyl-CoA Promiscuous trans-Acyltransferase. Chem. Biol. 2014, 21, 636–646.

(17) Passmore, M.; Gallo, A.; Lewandowski, J. R.; Jenner, M. Molecular Basis for Acyl Carrier Protein–Ketoreductase Interaction in Trans-Acyltransferase Polyketide Synthases. Chem. Sci. 2021, 12, 13676–13685.

(18) Fage, C. D.; Passmore, M.; Tatman, B. P.; Smith, H. G.; Jian, X.; Dissanayake, U. C.; Foran, M. E.; Cisneros, G. A.; Challis, G. L.; Lewandowski, J. R.; Jenner, M. Molecular Basis for Short-Chain Thioester Hydrolysis by Acyl Hydrolases in Trans-Acyltransferase Polyketide Synthases. JACS Au 2025, 5, 144–157.

(19) Mindrebo, J. T.; Misson, L. E.; Johnson, C.; Noel, J. P.; Burkart, M. D. Activity Mapping the Acyl Carrier Protein: Elongating Ketosynthase Interaction in Fatty Acid Biosynthesis. Biochemistry 2020, 59, 3626–3638.

(20) Miyanaga, A.; Iwasawa, S.; Shinohara, Y.; Kudo, F.; Eguchi, T. Structure-Based Analysis of the Molecular Interactions between Acyltransferase and Acyl Carrier Protein in Vicenistatin Biosynthesis. Proc. Natl. Acad. Sci. U.S.A. 2016, 113, 1802–1807.

(21) Miyanaga, A.; Ouchi, R.; Ishikawa, F.; Goto, E.; Tanabe, G.; Kudo, F.; Eguchi, T. Structural Basis of Protein-Protein Interactions between a trans-Acting Acyltransferase and Acyl Carrier Protein in Polyketide Disorazole Biosynthesis. J. Am. Chem. Soc. 2018, 140, 7970–7978.

(22) Misson, L. E.; Mindrebo, J. T.; Davis, T. D.; Patel, A.; Andrew McCammon, J.; Noel, J. P.; Burkart, M. D. Interfacial Plasticity Facilitates High Reaction Rate of E. Coli FAS Malonyl-CoA:ACP Transacylase, FabD. Proc. Natl. Acad. Sci. U.S.A., 2020, 117, 24224–24233.

(23) Honorato, R. V.; Trellet, M. E.; Jiménez-García, B.; Schaarschmidt, J. J.; Giulini, M.; Reys, V.; Koukos, P. I.; Rodrigues, J. P. G. L. M.; Karaca, E.; van Zundert, G. C. P.; Roel-Touris, J.; van Noort, C. W.; Jandová, Z.; Melquiond, A. S. J.; Bonvin, A. M. J. J. The HADDOCK2.4 Web Server for Integrative Modeling of Biomolecular Complexes. Nat. Protoc. 2024, 19, 3219–3241.

(24) Abramson, J.; Adler, J.; Dunger, J.; Evans, R.; Green, T.; Pritzel, A.; Ronneberger, O.; Willmore, L.; Ballard, A. J.; Bambrick, J.; Bodenstein, S. W.; Evans, D. A.; Hung, C.-C.; O’Neill, M.; Reiman, D.; Tunyasuvunakool, K.; Wu, Z.; Žemgulytė, A.; Arvaniti, E.; Beattie, C.; Bertolli, O.; Bridgland, A.; Cherepanov, A.; Congreve, M.; Cowen-Rivers, A. I.; Cowie, A.; Figurnov, M.; Fuchs, F. B.; Gladman, H.; Jain, R.; Khan, Y. A.; Low, C. M. R.; Perlin, K.; Potapenko, A.; Savy, P.; Singh, S.; Stecula, A.; Thillaisundaram, A.; Tong, C.; Yakneen, S.; Zhong, E. D.; Zielinski, M.; Žídek, A.; Bapst, V.; Kohli, P.; Jaderberg, M.; Hassabis, D.; Jumper, J. M. Accurate Structure Prediction of Biomolecular Interactions with AlphaFold 3. Nature 2024, 630, 493–500.

(25) Oefner, C.; Schulz, H.; D’Arcy, A.; Dale, G. E. Mapping the Active Site of Escherichia Coli Malonyl-CoA-Acyl Carrier Protein Transacylase (FabD) by Protein Crystallography. Acta Crystallogr. D Biol. Crystallogr. 2006, 62, 613–618.

(26) Liew, C. W.; Nilsson, M.; Chen, M. W.; Sun, H.; Cornvik, T.; Liang, Z.-X.; Lescar, J. Crystal Structure of the Acyltransferase Domain of the Iterative Polyketide Synthase in Enediyne Biosynthesis. J. Biol. Chem. 2012, 287, 23203– 23215.

(27) Ramírez-Palacios, C.; Wijma, H. J.; Thallmair, S.; Marrink, S. J.; Janssen, D. B. Computational Prediction of ω-Transaminase Specificity by a Combination of Docking and Molecular Dynamics Simulations. J. Chem. Inf. Model 2021, 61, 5569–5580.

(28) Kennedy, J.; Auclair, K.; Kendrew, S. G.; Park, C.; Vederas, J. C.; Hutchinson, C. R. Modulation of Polyketide Synthase Activity by Accessory Proteins during Lovastatin Biosynthesis. Science 1999, 284, 1368–1372.

(29) Meehan, M. J.; Xie, X.; Zhao, X.; Xu, W.; Tang, Y.; Dorrestein, P. C. FT-ICR-MS Characterization of Intermediates in the Biosynthesis of the α-Methylbutyrate Side Chain of Lovastatin by the 277 kDa Polyketide Synthase LovF. Biochemistry 2011, 50, 287–299.

(30) Ma, S. M.; Li, J. W. H.; Choi, J. W.; Zhou, H.; Lee, K. K. M.; Moorthie, V. A.; Xie, X.; Kealey, J. T.; Da Silva, N. A.; Vederas, J. C.; Tang, Y. Complete Reconstitution of a Highly Reducing Iterative Polyketide Synthase. Science 2009, 326, 589–592.

(31) Winter, J. M.; Sato, M.; Sugimoto, S.; Chiou, G.; Garg, N. K.; Tang, Y.; Watanabe, K. Identification and Characterization of the Chaetoviridin and Chaetomugilin Gene Cluster in Chaetomium Globosum Reveal Dual Functions of an Iterative Highly-Reducing Polyketide Synthase. J. Am. Chem. Soc. 2012, 134, 17900–17903.

(32) Qiao, K.; Chooi, Y.-H.; Tang, Y. Identification and Engineering of the Cytochalasin Gene Cluster from Aspergillus Clavatus NRRL 1. Metab. Eng. 2011, 13, 723–732.

(33) Winter, J. M.; Cascio, D.; Dietrich, D.; Sato, M.; Watanabe, K.; Sawaya, M. R.; Vederas, J. C.; Tang, Y. Biochemical and Structural Basis for Controlling Chemical Modularity in Fungal Polyketide Biosynthesis. J. Am. Chem. Soc. 2015, 137, 9885–9893.

