## Supplementary Information for "Molecular characterisation of the acyltransferase-acyl carrier protein interface in a fungal highly reducing polyketide synthase"

### 1. Supplementary Figures

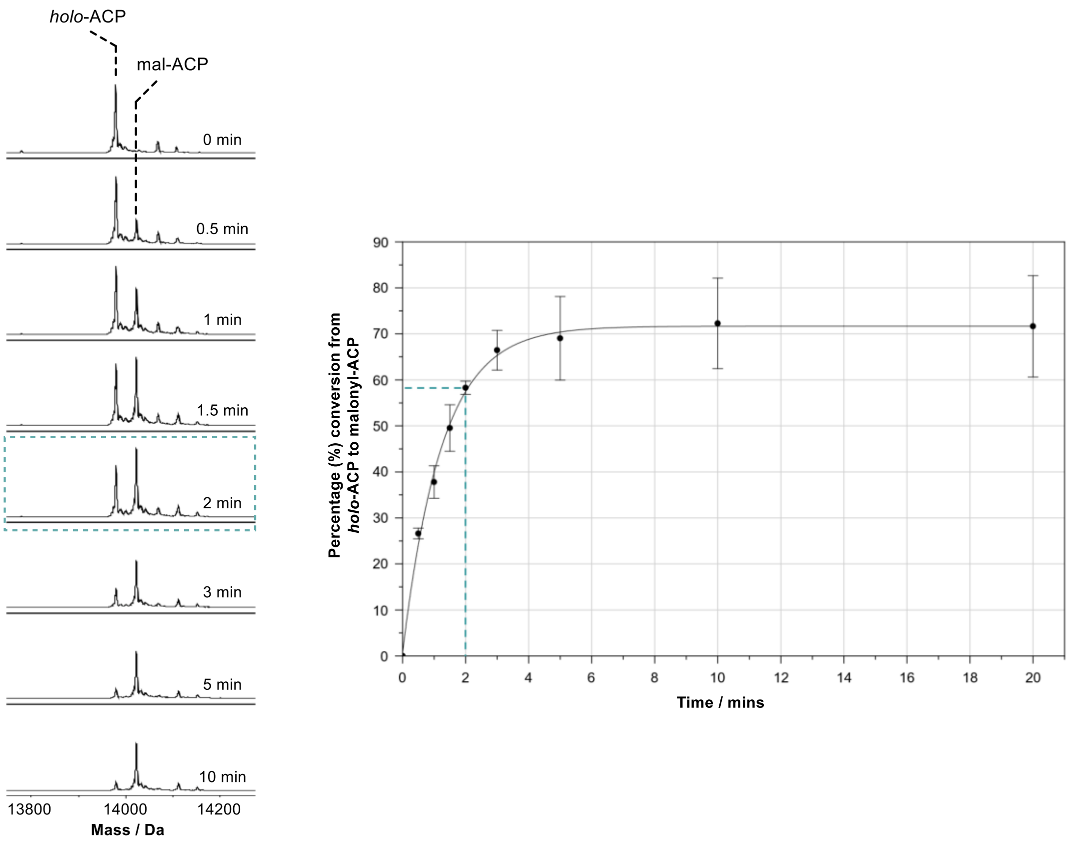


**Supplementary Figure S1 | Time-course of SimG AT-catalysed malonyl transfer to SimG ACP.** Stacked deconvoluted mass spectra (*left*) and plot (*right*) of the malonyl-transfer reaction between SimG AT and *holo*-SimG ACP domains over a 10 min time-course. The ratio between the *holo*-ACP and malonyl-ACP peaks was measured for each time point and plotted as percentage conversion. The 2-minute time point (indicated with a dashed line, 58.3 ± 1.44 %) was selected to assess the relative activities of the ACP domain X→Ala mutants, as it is situated in the linear region of the curve.

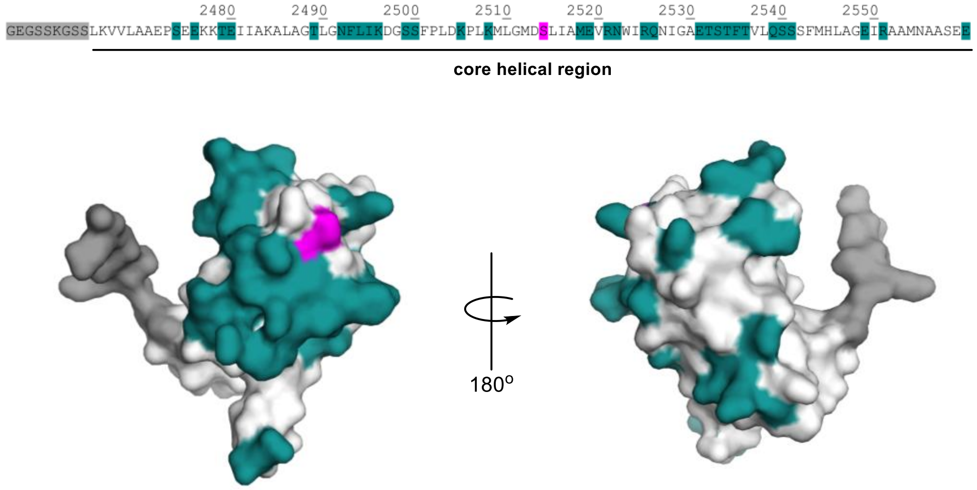


**Supplementary Figure S2 | Coverage of SimG ACP domain with X→Ala mutations.** Amino acid sequence (*top*) and AlphaFold 3 model (*bottom*) of the SimG ACP domain is shown. The core helical region is underlined and all X→Ala mutants are highlighted in teal. Approximately 49 % of the solvent exposed non-Ala/Gly residues from the helical core were covered by the scanning alanine mutagenesis approach. The Ppant attachment site, Ser2514, is highlighted in magenta.

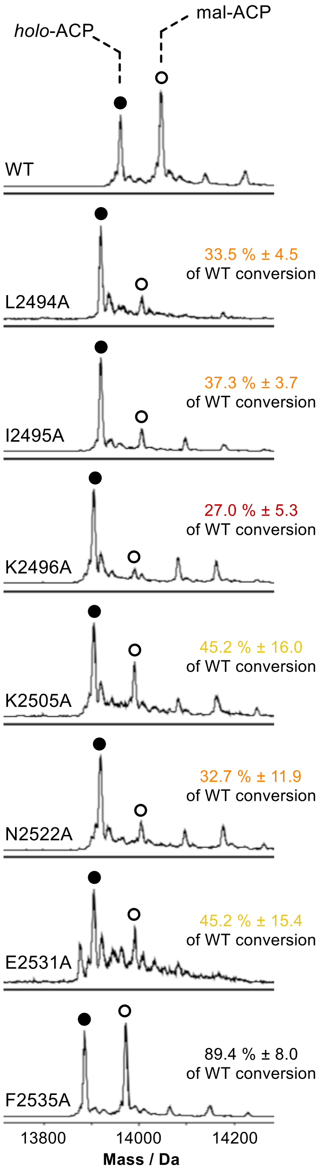


**Supplementary Figure S3 | Example spectra of malonyl transfer reactions for SimG ACP domain X→Ala mutants.** Deconvoluted mass spectra of SimG ACP domain (WT and selected X→Ala mutants) after 2 min malonyl transfer reaction with SimG AT domain. Values are reported relative to WT SimG ACP transfer (58.3 ± 1.44 %) and errors represent ± 1 standard deviation (1σ) from the mean, where $n$ = 3. Activities relative to WT are highlighted in red (< 30 %), orange (30 - 40 %), yellow (40 - 50 %), and black (> 50 %). The F2523A mutant is displayed as an example of a mutation that obtained near-WT levels of acylation.

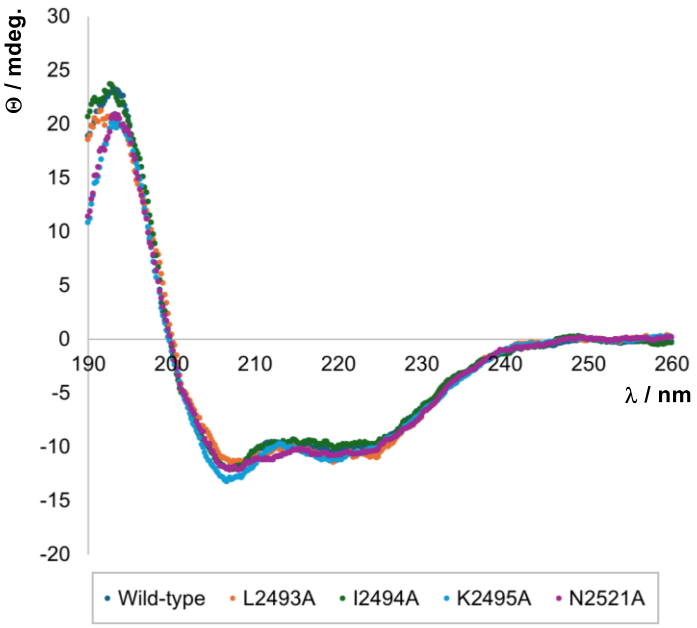


**Supplementary Figure S4 | Circular dichroism spectroscopic analysis of SimG ACP X→Ala mutants.** Overlaid CD spectra for wild-type SimG ACP domain and four alanine mutants which disrupted the interaction with SimG AT domain. The CD spectra obtained for each mutant are in agreement with the WT PksJ ACP4 domain, suggesting no loss / gain of secondary structure has occurred upon mutation.

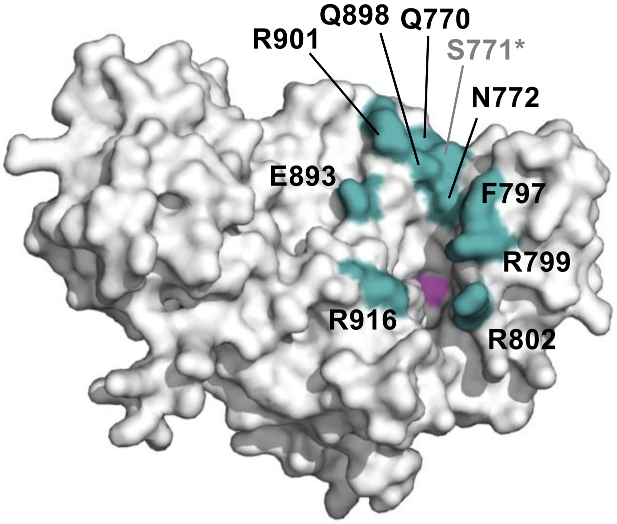


**Supplementary Figure S5 | SimG AT domain X→Ala mutations.** AlphaFold 3 model of SimG AT domain with residues surrounding the Ppant / CoA binding cleft highlighted in teal and labelled. All residues were mutated to Ala and yielded soluble protein. The S771A mutant showed dramatically decreased levels of acylation when incubated with malonyl-CoA (see **Supplementary Figure S6**), and was omitted from further assays. The catalytic Ser706 and His808 residues are highlighted in magenta.

a). b).

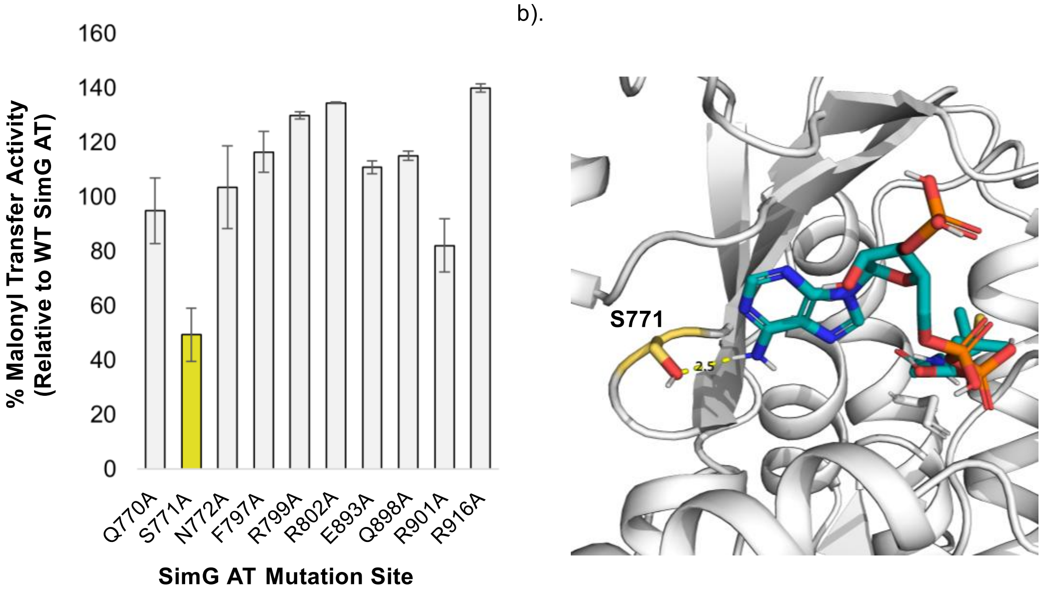


**Supplementary Figure S6 | Acylation of SimG AT domain X→Ala mutants with malonyl-CoA.** a). Bar chart showing percentage acylation of the X→Ala mutants generated for SimG AT domain, relative to wild-type SimG AT domain acylation. b). Alignment of the SimG AT domain AlphaFold 3 model with the *E. coli* FAS AT domain crystal structure with coenzyme A bound (PDB: 2G2Z). The proximity of Ser771 to the coenzyme A substrate likely provides a hydrogen bonding interaction which promotes acylation of the AT domain.

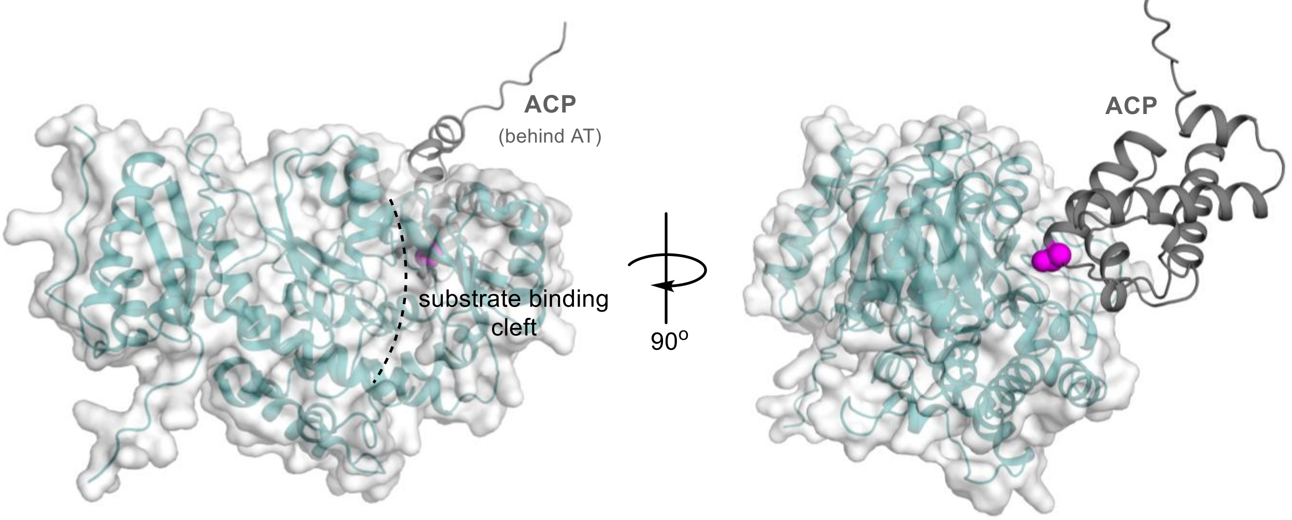


**Supplementary Figure S7 | AlphaFold of SimG AT:ACP complex.** Predicted of the SimG AT:ACP complex using AlphaFold. The ACP domain is placed behind the substrate binding cleft (indicated by a dashed line) of the AT domain. The site of Ppant attachment (Ser2514) is highlighted in magenta.

a). b).

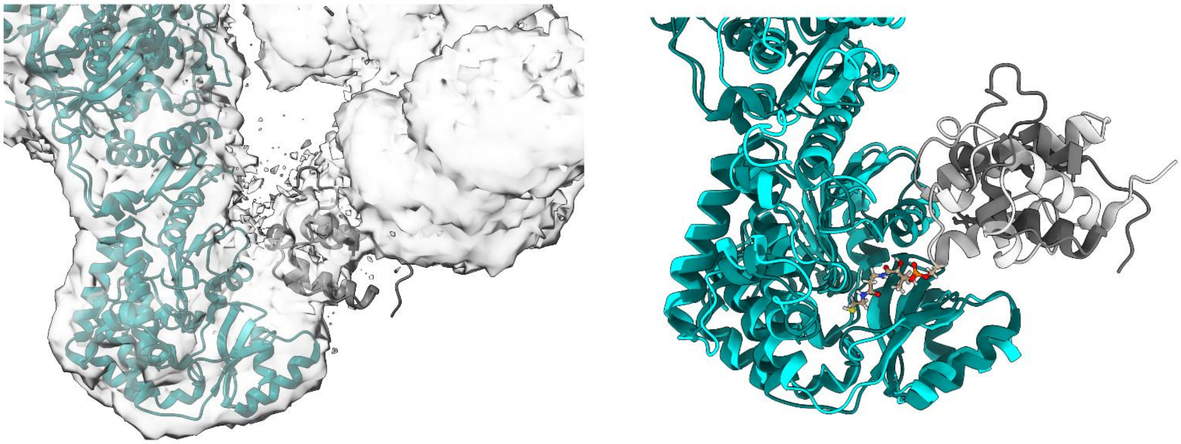

**Supplementary Figure S8 | Comparison of docked SimG AT:ACP complex with mFAS cryo-EM density map.** a). Overlay of the cryo-EM map (EMDB-43337) with atomic model of hFASN-NADPH state 1. MAT is indicated in teal. ACP fitted in partial density is indicated in dark grey. b). Overlay of the atomic model from a) with a representative conformation for the SimG AT-ACP complex determined in this work. AT is indicated in cyan. ACP is indicated in light grey. Post translationally modified Ser on ACP in the cryo-EM model is rendered with grey atoms. Loaded Ppant in our complex is also shown.

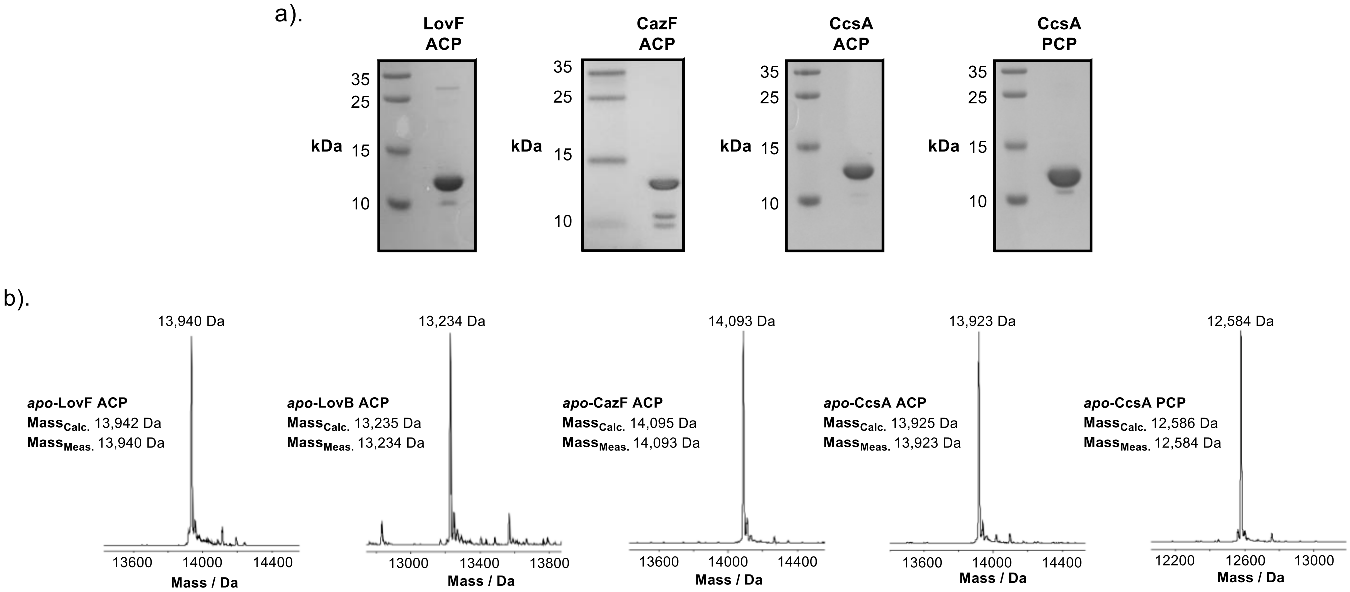


**Supplementary Figure S9 | SDS-PAGE and mass spectrometry analysis of purified recombinant proteins.** a). 15 % SDS-PAGE gel containing (from left to right): LovF ACP domain (~14 kDa), CazF ACP domain (~14 kDa), CcsA ACP domain (~14 kDa), and CcsA PCP domain (~13 kDa). b). Deconvoluted mass spectra of purified ACP / PCP domains with calculated and observed masses detailed. The mass spectrum of LovB ACP is shown as this is a modified construct from the previously reported construct, which contains additional His residues in the tag to improve purification.

a).

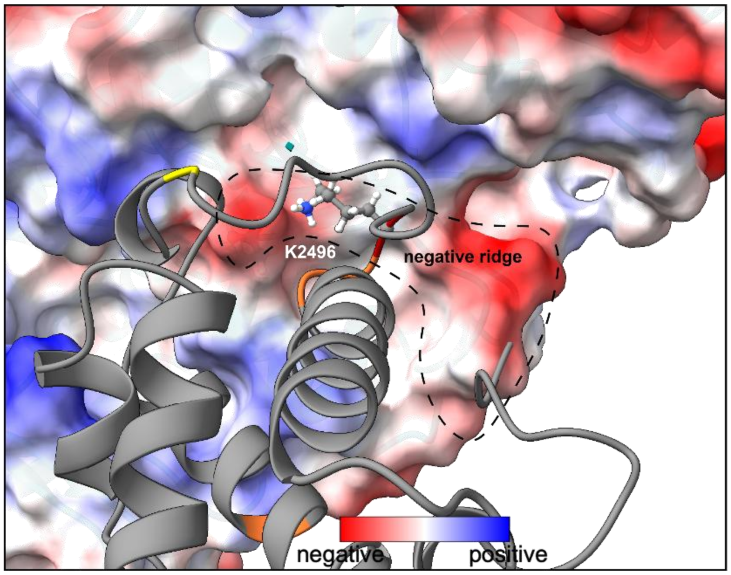


b).

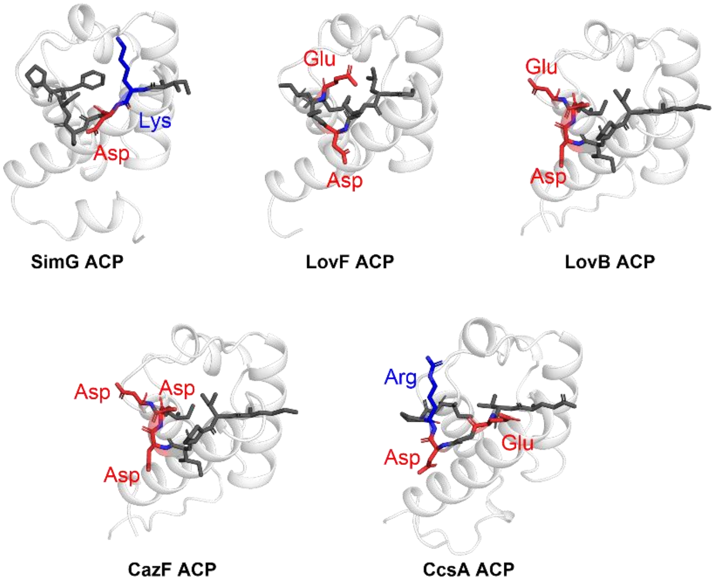


**Supplementary Figure S10 | Electrostatic interactions between SimG AT and cognate / non-cognate ACP domains.** a). Representative frame of the SimG ACP:AT complex following MD simulations (as shown in **Fig. 4a**). The surface electrostatics of the AT domain are displayed and the critical K2496 residue on the ACP domain is shown as sticks. The negatively charged ridge can be sampled by a positively charged residue (Lys / Arg) in the Loop I region. b). AlphaFold structural models of ACP domains used in malonyl transfer assays. The first 8 residues of the Loop I region are displayed as sticks, and negatively / positively charged residues are coloured red and blue, respectively. The presence of a positively charged residue in SimG ACP (cognate) and CcsA ACP (non-cognate) correlates with highest levels of malonyl transfer (**Fig. 5b**).

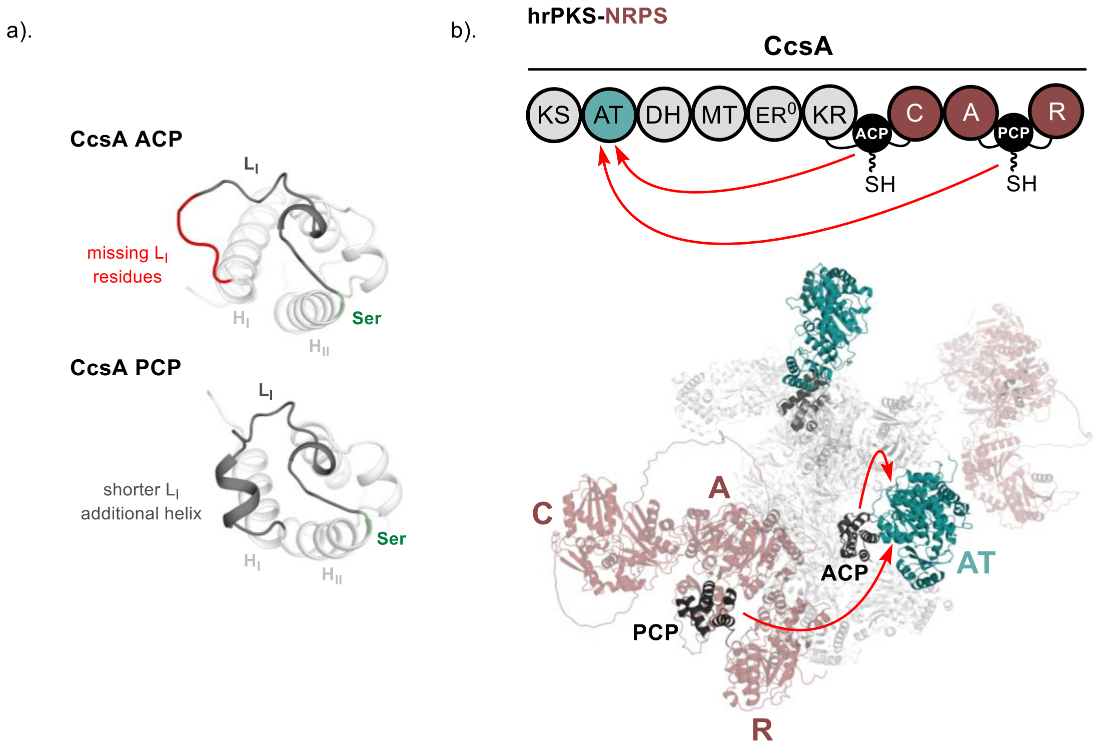


**Supplementary Figure S11 | Structural differences and accessibility of carrier protein domains in PKS-NRPS hybrids.** a). Structural comparison of AlphaFold models for CcsA ACP (*top*) and PCP (*bottom*) domains. The $L_I$ region is highlighted in dark grey, and the residues missing from CcsA PCP are indicated in red. The site of Ppant attachment is highlighted in green. b). Domain arrangement (*top*) and AlphaFold model (*bottom*) of the entire CcsA protein. The protein was assembled into a dimer by structural alignment of two copies with the LovB hrPKS (PDB: 7CPX).[1] Whilst the flexible linkers may allow the PCP domain to access the AT domain, a combination of improved transacylation for the ACP domain (**Fig. 5b**), and engagement with the A, C and R domains of the NRPS region, likely mean that the PCP domain is not loaded with malonyl during catalysis.

### 2. Methods

#### 2.1. Molecular cloning and mutagenesis.

The cloning of SimG AT and SimG ACP plasmids has been reported previously.[2] The plasmids for LovF ACP, LovB ACP, CazF ACP, CcsA ACP and CcsA PCP were gene synthesised (Epoch Life Sciences) and subcloned into pET24a with a custom $pHis_8$ tag (see **Section 3.1**). All SimG ACP and AT mutants were constructed using the Q5 site directed mutagenesis kit (NEB), using primers detailed in **Tables S1** and **S3** introducing a X→Ala mutation in each case. The resulting PCR products were processed according to the manufactures protocol, and resulting plasmids sequenced to verify correct mutation.

#### 2.2. Protein overexpression and purification.

Overexpression and purification of SimG AT and ACP domains has been reported previously.[2] The same procedure was applied to all SimG ACP X→Ala mutants and non-cognate ACP domains (LovF ACP, LovB ACP, CazF ACP, CcsA ACP and CcsA PCP) used in this study. Briefly, an aliquot of chemically competent *E. coli* BL21 Star (DE3) cells (50 µL) was transformed with relevant plasmid DNA. Transformed cells were plated onto LB agar plates containing kanamycin (50 µg/mL) and incubated overnight at 37˚C. A single colony was picked and used to inoculate LB media (10 mL) containing kanamycin (50 µg/mL) and incubated overnight at 37˚C with shaking (180 rpm). The overnight culture was used to inoculate a flask of LB media (1 L) with kanamycin (50 µg/mL) which was left to grow at 37˚C with shaking (180 rpm) until an $OD_{600}$ of 0.7 - 1.0 was reached. Protein expression was induced by the addition of IPTG (250 µM) and incubated at 15 ˚C with shaking (180 rpm) overnight.

Cells were centrifuged (4000 rpm, 15 minutes, 4 ˚C) and resuspended in loading buffer (20 mM Tris HCl, 100 mM NaCl, 20 mM imidazole, pH 8.0). The cells were lysed by high pressure (20 psi) cell disruption and centrifuged (17,000 rpm, 30 minutes, 4˚C) to pellet the insoluble cell components. The cell lysate was filtered through a 0.45 µm syringe filter and loaded onto a 1 mL HiTrap™ Nickel FastFlow column. Following a wash with loading buffer (10 mL), the protein was eluted from the column with buffers of increasing imidazole concentrations (5 mL of 50 mM, 3 mL of 100 mM, 3 mL of 200 mM, 3 mL of 300 mM). Fractions were checked for the protein of interest by SDS-PAGE and those containing protein were concentrated and exchanged into storage buffer (20 mM Tris HCl, 100 mM NaCl, pH 7.6) using an appropriate size Vivaspin centrifugal concentrator (4000 rpm, 4 ˚C). Once concentrated to approx. 1 mL, 50 µL aliquots of protein were flash frozen in liquid $N_2$ and stored at -80 ˚C.

#### 2.3. Conversion of *apo*- to *holo*-ACP / PCP domains.

All *apo*-ACP / PCP (200 µM) species (including X→Ala mutants) were converted to their *holo*-ACP / PCP form by incubation in storage buffer (20 mM Tris base, 100 mM NaCl, pH 7.6) supplemented with 10 mM $MgCl_2$, 2 µM Sfp, and 1 mM CoA in a total of 50 µL for 30 minutes at 25 ˚C. Samples were diluted 10-fold with milliQ $H_2O$ and analysed by UHPLC-ESI-Q-TOF-MS for successful conversion.

#### 2.4. SimG AT transacylation time-course.

SimG *holo*-ACP domain (50 µM) was incubated with SimG AT domain (1 µM) and malonyl-CoA (250 µM) in storage buffer (20 mM Tris base, 100 mM NaCl, pH 7.6) made up to a final volume of a final volume of 100 µL at 25 ºC. The reaction was quenched with formic acid (1 % v/v final) at intervals between 0.5 → 20 mins following AT domain incubation, before dilution with milliQ $H_2O$ and analysis by UHPLC-ESI-Q-TOF-MS. Raw mass spectra were subjected to deconvolution (Bruker MaxEnt), taking an average of all charge states. Using the raw peak intensities for *holo*-ACP and acyl-ACP, percentage conversion at a given time point was calculated using the following equation:

$$\%\ acyl\text{-ACP} = \frac{Inten._{acyl\text{-}ACP}}{(Inten._{holo\text{-ACP}} + Inten._{\text{acyl-ACP}})}$$

All time-point measurements were conducted in triplicate and average values were plotted. The average acylation value at the 2 min time point was 58.3 ± 1.44 %.

### 2.5. Assessment of transacylation activity for SimG ACP X→Ala mutant library.

Each SimG ACP domain X→Ala mutant (50 µM) was incubated with SimG AT domain (1 µM) and malonyl-CoA (250 µM) in storage buffer (20 mM Tris base, 100 mM NaCl, pH 7.6) made up to a final volume of a final volume of 50 µL at 25 ºC. The reaction was quenched with formic acid (1 % v/v final) after 2 min, and samples were diluted 5-fold with milliQ $H_2O$ for analysis by UHPLC-ESI-Q-TOF-MS. Percentage conversion to the malonyl-ACP domain species for each mutant was calculated as outlined in **Section 2.4**. This conversion was made relative to the wild-type ACP conversion using the following equation:

$$SimG\ ACP\ (X \rightarrow Ala)\ activity = \left(\frac{\%\ acyl\text{-ACP (X→Ala } mutant\text{)}}{58.3\ (\text{WT } activity)}\right) \times 100$$

### 2.6. UHPLC-ESI-Q-TOF-MS analysis of SimG ACP.

All transacylation assays were analysed on a Bruker MaXis II ESI-Q-TOF-MS connected to a Dionex 3000 RS UHPLC fitted with an ACE C4-300 RP column (100 x 2.1 mm, 5 µm, 30 °C). The column was eluted with a linear gradient of 5 – 100% MeCN containing 0.1 % formic acid over 30 min. The mass spectrometer was operated in positive ion mode with a scan range of 200 – 3000 *m/z*. Source conditions were: end plate offset at −500 V; capillary at −4500 V; nebulizer gas ($N_2$) at 1.8 bar; dry gas ($N_2$) at 9.0 L $min^{-1}$ ; dry temperature at 200 °C. Ion transfer conditions were: ion funnel RF at 400 Vpp; multiple RF at 200 Vpp; quadrupole low mass at 200 *m/z*; collision RF at 2000 Vpp; transfer time at 110.0 µs; pre-pulse storage time at 10.0 µs. Measured masses for the intact *apo*-ACP and AT species are displayed in **Table S2** and **S4**.

### 2.7. Circular dichroism analyses of SimG ACP and X→Ala mutants.

Circular dichroism spectra of SimG ACP and associated mutants were acquired on a JASCO J-1500 spectropolarimeter, using a 1 mm path length quartz cuvette. Spectra were acquired between 190 and 260 nm at room temperature. Before analysis protein samples were exchanged into CD buffer (0.5 mM Tris HCl, 5 mM NaCl, pH 7.4) and adjusted to a final concentration of 0.1 mg/mL.

### 2.8. AlphaFold 3 model generation, residue-guided protein docking and molecular dynamics.

#### *2.8.1. AlphaFold 3 model generation of SimG AT and SimG ACP domains.*

Structural models of SimG AT and ACP domains were generated using AlphaFold 3.[3] Input sequences for each domain did not include His-Tag / non-native amino acids.

#### *2.8.2. Residue-guided protein docking of the SimG AT:ACP complex.*

Docking simulations between the SimG ACP and AT domain were performed using the High Ambiguity Driven DOCKing (HADDOCK 2.4) webserver.[4] Simulations were conducted using structural models of each domain, as outlined in **Section 2.8.1**. Residues identified from alanine scanning experiments were defined as 'active' in the docking simulation, and Ser2514 on the ACP domain defined as 'passive' as it is the Ppant attachment site. Results from the docking were evaluated for shape / charge complementarity and positioning of Ser2514 with respect to the substrate binding cleft. A structure from the highest-scoring docking solution cluster was taken forward for molecular dynamics.

#### *2.8.3. Molecular dynamic simulations of SimG holo-ACP:malonyl-AT complex.*

The co-ordinates for simulation of the SimG AT:ACP complex were taken from the first model of the highest scoring cluster from the docking simulation (see **Section 2.8.2**). A malonate was manually attached to Ser2514 using previous malonyl-AT structures as a guide[5,6], and the Ppant arm was placed manually in virtual reality using ChimeraX.[7–9] Molecular dynamics (MD) simulations were performed using the AMBER ff19SB force field.[10] Ppant and *O*-malonyl-serine were parametrised using ANTECHAMBER[11] and GAFF2 force field with ABCG2 charge model.[12] The resulting structure was neutralised with $Na^+$ ions and solvated with OPCBOX water, such that no atom belonging to the complex was less than 10 Å from any box edge, using the LEaP module.[13] Additional $Na^+$ and $Cl^-$ ions were added to obtain 100 final salt concentration. MD heating, equilibration, and production steps were performed using the GPU accelerated AMBER24 software[14] on an HPC cluster equipped with Nvidia L40 graphics cards. Simulation used the SHAKE algorithm[15] to constrain all protein bonds involving a hydrogen atom; a 2.0 fs time-step was used in these simulations. Long-range electrostatics were calculated using the Particle Mesh Ewald (PME) method with a 12.0 Å cut-off.[16] PME was used for nonbonded interactions. In all simulations, the Langevin thermostat ($\gamma$ = 2.0 ps$^{-1}$) was used to maintain temperature control.[17] The solvated protein was then equilibrated by carrying out a short minimization, 50 ps of heating and 50 ps of density equilibration with weak restraints on the protein followed by 500 ps of constant pressure equilibration at 300 K. After a two-step minimization process, in which solvent molecules were allowed to relax before the entire system was minimized, the system was slowly heated to 300 K over 0.1 ns in a canonical ensemble (NVT) simulation, then equilibrated for 2 ns by performing isothermal-isobaric (NPT) simulations at 300K using a Berendsen barostat.[18] Three independent repeats of 250 ns production using classic approach (with no acceleration) run were performed with simulation frames written every 20 ps for analysis. Additional 200 ns accelerated MD (aMD) was run from a restart point of the third repeat of the classical MD. For aMD frames were written every 20 ps. The aMD modification of the potential were defined as:

$$V(r)^* = V(r) + \Delta V(r)$$

$$\Delta V(r) = \frac{(E_P - V(r))^2}{(\alpha_P + E_P - V(r)} + \frac{(E_D - V(r))^2}{(\alpha_D + E_D - V_D(r)}$$

where $V(r)$ is the normal potential and $V_D(r)$ is the normal torsion potential, with $E_p$ = -400203.9002 kcal mol$^{-1}$; $\alpha_p$ = 27345.8 mol$^{-1}$; $E_D$ = 4831.4334 kcal mol$^{-1}$; $\alpha_D$ = 385.7 kcal mol$^{-1}$ (estimated from the cMD simulation, see Amber manual). For all the classic and accelerated simulations, long-range electrostatics were calculated using the PME method with 8.0 Å and 10.0 Å cutoffs, respectively. A combination of CCPTRAJ[19], Chimera[20] and ChimeraX[21] were used for analysis of the trajectories. ChimeraX was used throughout the course to prepare and visualize the structures.

The repeats of 250 ns cMD trajectories and one repeat of the 200 ns aMD trajectory stripped of water and ions with frames every 20 ns are available for download at https://doi.org/10.5281/zenodo.17272938.

### 3. Sequences, Tables and Accession Numbers

#### 3.1. Amino acid sequences

Amino acid sequences for SimG hrPKS, SimG AT domain and SimG ACP domain have been reported previously.[2]

**pHis$_8$_LovF ACP** (*Aspergillus terreus*, Q9Y7D5)

```
MKHHHHHHHH GGLVPRGSHG S-
NHGALSLTPA EDDNLHARLN RAISQQESIA VIMEAMSCKL ISMFGLTDSE MSATQTLAGI
GVDSLVAIEL RNWITAKFNV DISVFELMEG RTIAKVAEVV LQRYKA-
```

MW (including His-tag) = 13,941.94 Da
Ppant attachment site highlighted in Cyan.

**pHis$_8$_LovB ACP*** (*Aspergillus terreus*, Q9Y7D5)

```
MKHHHHHHHH GGLVPRGSHG S-
GEGAAGSKGS VKEQLLQATN LDQVRQIVID GLSAKLQVTL QIPDGESVHP TIPLIDQGVD
SLGAVTVGTW FSKQLYLDLP LLKVLGGASI TDLANEAAAR LPPSS-
```

MW (including His-tag) = 13,234.99 Da
Ppant attachment site highlighted in Cyan.
* core amino acid sequence based on previously published LovB ACP construct.[22]

**pHis$_8$_CazF ACP** (*Chaetomium globosum*, A0A0K0MCJ4)

```
MKHHHHHHHH GGYVPRGSHG S-
GAQAAGQQLG DADVKPLSAQ LQEAGSRDEA TRLVGDAIAS KLGDIFMIPI DDIDLAKSPA
LHGVDSLVAV ELRNMLMLQA AADISIFSIM QSASLGALAS DVVAKSSHVE IAAGA
```

MW (including His-tag) = 14,094.91 Da
Ppant attachment site highlighted in Cyan.

**pHis$_8$_CcsA ACP** (*Aspergillus clavatus*, A1CLY8)

```
MKHHHHHHHH GGLVPRGSHG S-
STSAAPVKVR LAEASSSADI YDIISDAFVT KLKTSLQVEG DRPIVDLTAD TLGIDSLVAV
DIRSWFIKEL QVEIPVLKIL SGATVGEMVT QAQELLPKEL TPNLDPNA
```

MW (including His-tag) = 13,924.85 Da
Ppant attachment site highlighted in Cyan.

**pHis$_8$_CcsA PCP** (*Aspergillus clavatus*, A1CLY8)

```
MKHHHHHHHH GGLVPRGSHG S-
ADTGTDESPS MAKMRDVWAT VIPQEVLAHF ELGPASNFFQ VGGDSMLLVR LQTEINKVFG
TSISLFQLFD ASSLTGMVSL IDHSESTSQR SEVD
```

MW (including His-tag) = 12,586.04 Da
Ppant attachment site highlighted in Cyan.

### 3.2. Supplementary Tables

**Table S1.** Primers for SimG ACP domain X →Ala mutants.

| SimG ACP Mutant | Primers For / Rev (5'→3') | Annealing Temperature (°C) |
|---|---|---|
| S2474A | Forward: CGCCGAGCCAGCAGAGGAGAAGAA<br>Reverse: GCCAGGACGACCTTGAGC | 69 |
| E2476A | Forward: CGAGCCATCTGCGGAGAAGAAGACC<br>Reverse: GCGGCCAGGACGACCTTG | 72 |
| T2479A | Forward: GGAGAAGAAGGCGGAGATCATCGCCAAGGC<br>Reverse: TCAGATGGCTCGGCGGCC | 72 |
| E2480A | Forward: GAAGAAGACCGCGATCATCGCCAAG<br>Reverse: TCCTCAGATGGCTCGGCG | 67 |
| T2489A | Forward: CCTCGCCGGCGCACTCGGGAACT<br>Reverse: GCCTTGGCGATGATCTCGG | 69 |
| N2492A | Forward: CACTCTCGGGGCGTTCCTCATCAAAGACGGGAGCAGCTTCCC<br>Reverse: CCGGCGAGGGCCTTGGCG | 72 |
| F2493A | Forward: TCTCGGGAACGCGCTCATCAAAGACGGGAGCAGCTTC<br>Reverse: GTGCCGGCGAGGGCCTTG | 72 |
| L2494A | Forward: CGGGAACTTCGCGATCAAAGACGGGAGCAGCTTCCC<br>Reverse: AGAGTGCCGGCGAGGGCC | 72 |
| I2495A | Forward: GAACTTCCTCGCGAAAGACGGGAGCAGCTTCCCAC<br>Reverse: CCGAGAGTGCCGGCGAGG | 71 |
| K2496A | Forward: CTTCCTCATCGCGGACGGGAGCAGCTTCCCAC<br>Reverse: TTCCCGAGAGTGCCGGCG | 70 |
| S2499A | Forward: CAAAGACGGGGCGAGCTTCCCACTC<br>Reverse: ATGAGGAAGTTCCCGAGAG | 60 |
| S2500A | Forward: AGACGGGAGCGCGTTCCCACTCG<br>Reverse: TTGATGAGGAAGTTCCCG | 60 |
| K2505A | Forward: CCCACTCGACGCACCCCTCAAGATG<br>Reverse: AAGCTGCTCCCGTCTTTG | 62 |
| K2508A | Forward: CAAGCCCCTCGCAATGCTCGGCATG<br>Reverse: TCGAGTGGGAAGCTGCTC | 66 |
| M2518A | Forward: TTTGATCGCCGCAGAGGTGCGGAAC<br>Reverse: GAATCCATGCCGAGCATC | 62 |
| E2519A | Forward: GATCGCCATGGCAGTGCGGAACT<br>Reverse: AAAGAATCCATGCCGAGC | 63 |
| R2521A | Forward: CATGGAGGTGGCAAACTGGATCC<br>Reverse: GCGATCAAAGAATCCATG | 57 |
| N2522A | Forward: GGAGGTGCGGGCATGGATCCGAC<br>Reverse: ATGGCGATCAAAGAATCC | 60 |
| R2525A | Forward: GAACTGGATCGCACAGAACATCGGCGC<br>Reverse: CGCACCTCCATGGCGATC | 67 |
| Q2526A | Forward: CTGGATCCGAGCAAACATCGGCGCCGAGACAAG<br>Reverse: TTCCGCACCTCCATGGCG | 68 |
| E2531A | Forward: CATCGGCGCCGCAACAAGCACGT<br>Reverse: TTCTGTCGGATCCAGTTCCG | 67 |
| T2532A | Forward: CGGCGCCGAGGCAAGCACGTTCA<br>Reverse: ATGTTCTGTCGGATCCAGTTCCGCAC | 72 |
| S2533A | Forward: CGCCGAGACAGCAACGTTCACCG<br>Reverse: CCGATGTTCTGTCGGATC | 60 |
| T2534A | Forward: CGAGACAAGCGCATTCACCGTGC<br>Reverse: GCGCCGATGTTCTGTCGG | 67 |
| F2535A | Forward: GACAAGCACGGCAACCGTGCTACAGAG<br>Reverse: TCGGCGCCGATGTTCTGT | 66 |
| T2536A | Forward: AAGCACGTTCGCAGTGCTACAGA<br>Reverse: GTCTCGGCGCCGATGTTC | 65 |
| Q2539A | Forward: CACCGTGCTAGCAAGCTCTTCCTTCATGCATCTG<br>Reverse: AACGTGCTTGTCTCGGCG | 65 |
| S2540A | Forward: CGTGCTACAGGCATCTTCCTTCATGCATC<br>Reverse: GTGAACGTGCTTGTCTCG | 60 |
| S2541A | Forward: GCTACAGAGCGCATCCTTCATGC<br>Reverse: ACGGTGAACGTGCTTGTC | 63 |
| E2549A | Forward: TCTGGCCGGCGCAATCAGGGCGG<br>Reverse: TGCATGAAGGAAGAGCTCTGTAGCAC | 72 |
| R2551A | Forward: CGGCGAGATCGCAGCGGCGATGA<br>Reverse: GCCAGATGCATGAAGGAAG | 64 |
| E2560A | Forward: GGCGAGCGAGGCATGAAAGCTTGCGG<br>Reverse: GCATTCATCGCCGCCCTG | 68 |

**Table S2.** Measured masses of SimG ACP domain X →Ala mutants.

| SimG ACP Mutant | Calculated Mass (Da) | Measured Mass (Da) |
|---|---|---|
| S2474A | 13607.56 | 13607.50 |
| E2476A | 13565.52 | 13564.60 |
| T2479A | 13593.53 | 13591.73 |
| E2480A | 13565.52 | 13564.85 |
| T2489A | 13593.53 | 13593.56 |
| N2492A | 13580.53 | 13579.48 |
| F2493A | 13547.46 | 13547.45 |
| L2494A | 13581.48 | 13580.63 |
| I2495A | 13581.48 | 13580.81 |
| K2496A | 13566.46 | 13565.62 |
| S2499A | 13607.56 | 13606.69 |
| S2500A | 13607.56 | 13606.60 |
| K2505A | 13566.46 | 13566.56 |
| K2508A | 13566.46 | 13565.88 |
| M2518A | 13563.45 | 13562.63 |
| E2519A | 13565.52 | 13564.00 |
| R2521A | 13538.45 | 13537.69 |
| N2522A | 13580.53 | 13579.88 |
| R2525A | 13538.45 | 13537.69 |
| Q2526A | 13566.51 | 13565.63 |
| E2531A | 13565.52 | 13564.69 |
| T2532A | 13593.53 | 13592.63 |
| S2533A | 13607.56 | 13606.88 |
| T2534A | 13593.53 | 13592.81 |
| F2535A | 13547.46 | 13546.69 |
| T2536A | 13593.53 | 13592.63 |
| Q2539A | 13566.51 | 13566.56 |
| S2540A | 13607.56 | 13605.65 |
| S2541A | 13607.56 | 13606.69 |
| E2549A | 13565.52 | 13564.56 |
| R2551A | 13538.45 | 13537.69 |
| E2560A | 13565.52 | 13564.69 |

**Table S3.** Primers for SimG AT domain X →Ala mutants

| SimG AT Mutant | Primers For / Rev (5'→3') | Annealing Temperature (°C) |
|---|---|---|
| Q770A | Forward: CGAGAACAGTGCAAGCAACGTCACGCTGTCGG<br>Reverse: CAGGCGATGTTGACCCCC | 66 |
| S771A | Forward: GAACAGTCAGGCAAACGTCACGCTGTCGG<br>Reverse: TCGCAGGCGATGTTGACC | 64 |
| N772A | Forward: CAGTCAGAGCGCAGTCACGCTGTCG<br>Reverse: TTCTCGAGGCGATGTTG | 61 |
| F797A | Forward: ACCGGGGGTCGCAGCTCGTTTGC<br>Reverse: CTCTCGGCCTTGAGCGTC | 65 |
| R799A | Forward: GGTCTTTGCTGCATTGCTCCGGGTAGAGAAGGC<br>Reverse: CCCGGTCTCTCGGCCTTG | 67 |
| R802A | Forward: TCGTTTGCTCGCAGTAGAGAAGGCGTATCACTC<br>Reverse: GCAAAGACCCCCGGTCTC | 66 |
| E893A | Forward: TCCCGCCCTTGCAGGACCAGTCG<br>Reverse: TGAGGTCCCAACTCGATTAGGAC | 69 |
| Q898A | Forward: ACCAGTCGGCGCAATTCTCCGAGACCTGG<br>Reverse: CCTTCAAGGGCGGGATGA | 65 |
| R901A | Forward: CCAGATTCTCGCAGACCTGGGTCG<br>Reverse: CCGACTGGTCCTTCAAGG | 63 |
| R916A | Forward: GACCTGTATCGCAGGCAGCGAGTG<br>Reverse: CCAAGGTAGATGTCGTCTG | 60 |

**Table S4.** Measured masses of SimG AT domain X →Ala mutants.

| SimG AT Mutant | Calculated Mass (Da) | Measured Mass (Da) |
|---|---|---|
| Q770A | 51104.67 | 51101.44 |
| S771A | 51145.67 | 51142.50 |
| N772A | 51118.70 | 51117.38 |
| F797A | 51085.62 | 51083.59 |
| R799A | 51076.61 | 51074.66 |
| R802A | 51076.61 | 51074.80 |
| E893A | 51103.68 | 51100.53 |
| Q898A | 51104.67 | 51101.84 |
| R901A | 51076.61 | 51073.39 |
| R916A | 51076.61 | 49075.00* |

* degradation observed, but full activity in assays.

### 4. References


(1) Wang, J.; Liang, J.; Chen, L.; Zhang, W.; Kong, L.; Peng, C.; Su, C.; Tang, Y.; Deng, Z.; Wang, Z. Structural Basis for the Biosynthesis of Lovastatin. *Nat. Commun.* **2021**, *12*, 867.

(2) Foran, M. E.; Auckloo, N. B.; Ho, Y. T. C.; Liu, S.; Hai, Y.; Jenner, M. Biochemical Dissection of a Fungal Highly Reducing Polyketide Synthase Condensing Region Reveals Basis for Acyl Group Selection. *Chem. Sci.* **2025**, *16*, 13173–13182.

(3) Abramson, J.; Adler, J.; Dunger, J.; Evans, R.; Green, T.; Pritzel, A.; Ronneberger, O.; Willmore, L.; Ballard, A. J.; Bambrick, J.; Bodenstein, S. W.; Evans, D. A.; Hung, C.-C.; O'Neill, M.; Reiman, D.; Tunyasuvunakool, K.; Wu, Z.; Žemgulytė, A.; Arvaniti, E.; Beattie, C.; Bertolli, O.; Bridgland, A.; Cherepanov, A.; Congreve, M.; Cowen-Rivers, A. I.; Cowie, A.; Figurnov, M.; Fuchs, F. B.; Gladman, H.; Jain, R.; Khan, Y. A.; Low, C. M. R.; Perlin, K.; Potapenko, A.; Savy, P.; Singh, S.; Stecula, A.; Thillaisundaram, A.; Tong, C.; Yakneen, S.; Zhong, E. D.; Zielinski, M.; Žídek, A.; Bapst, V.; Kohli, P.; Jaderberg, M.; Hassabis, D.; Jumper, J. M. Accurate Structure Prediction of Biomolecular Interactions with AlphaFold 3. *Nature* **2024**, *630*, 493–500.

(4) Honorato, R. V.; Trellet, M. E.; Jiménez-García, B.; Schaarschmidt, J. J.; Giulini, M.; Reys, V.; Koukos, P. I.; Rodrigues, J. P. G. L. M.; Karaca, E.; van Zundert, G. C. P.; Roel-Touris, J.; van Noort, C. W.; Jandová, Z.; Melquiond, A. S. J.; Bonvin, A. M. J. J. The HADDOCK2.4 Web Server for Integrative Modeling of Biomolecular Complexes. *Nat. Protoc.* **2024**, *19*, 3219–3241.

(5) Oefner, C.; Schulz, H.; D'Arcy, A.; Dale, G. E. Mapping the Active Site of Escherichia Coli Malonyl-CoA-Acyl Carrier Protein Transacylase (FabD) by Protein Crystallography. *Acta Crystallogr. D Biol. Crystallogr.* **2006**, *62*, 613–618.

(6) Liew, C. W.; Nilsson, M.; Chen, M. W.; Sun, H.; Cornvik, T.; Liang, Z.-X.; Lescar, J. Crystal Structure of the Acyltransferase Domain of the Iterative Polyketide Synthase in Enediyne Biosynthesis. *J. Biol. Chem.* **2012**, *287*, 23203–23215.

(7) Pettersen, E. F.; Goddard, T. D.; Huang, C. C.; Meng, E. C.; Couch, G. S.; Croll, T. I.; Morris, J. H.; Ferrin, T. E. UCSF ChimeraX: Structure Visualization for Researchers, Educators, and Developers. *Protein Sci* **2021**, *30*, 70–82.

(8) Goddard, T. D.; Huang, C. C.; Meng, E. C.; Pettersen, E. F.; Couch, G. S.; Morris, J. H.; Ferrin, T. E. UCSF ChimeraX: Meeting Modern Challenges in Visualization and Analysis. *Protein Sci.* **2018**, *27*, 14–25.

(9) Goddard, T. D.; Brilliant, A. A.; Skillman, T. L.; Vergenz, S.; Tyrwhitt-Drake, J.; Meng, E. C.; Ferrin, T. E. Molecular Visualization on the Holodeck. *J. Mol. Biol.* **2018**, *430*, 3982–3996.

(10) Tian, C.; Kasavajhala, K.; Belfon, K. A. A.; Raguette, L.; Huang, H.; Migues, A. N.; Bickel, J.; Wang, Y.; Pincay, J.; Wu, Q.; Simmerling, C. Ff19SB: Amino-Acid-Specific Protein Backbone Parameters Trained against Quantum Mechanics Energy Surfaces in Solution. *J. Chem. Theory Comput.* **2020**, *16*, 528–552.

(11) Wang, J.; Wang, W.; Kollman, P. A.; Case, D. A. Automatic Atom Type and Bond Type Perception in Molecular Mechanical Calculations. *J. Mol. Graph Model* **2006**, *25*, 247–260.

(12) He, X.; Man, V. H.; Yang, W.; Lee, T.-S.; Wang, J. A Fast and High-Quality Charge Model for the next Generation General AMBER Force Field. *J. Chem. Phys.* **2020**, *153*, 114502.

(13) Schafmeister, C. E. A. F.; Ross, W. S.; Romanovski, V. LEap. *University of California, San Francisco* **1995**.

(14) Case, D. A.; Aktulga, H. M.; Belfon, K.; Ben-Shalom, I. Y.; Berryman, J. T.; Brozell, S. R.; Carvahol, F. S.; Cerutti, D. S.; Cheatham, T. E.; Cisneros, G. A.; Cruzeiro, V. W. D.; Darden, T. A.; Forouzesh, N.; Ghazimirsaeed, M.; Giambaşu, G.; Giese, T.; Gilson, M. K.; Gohlke, H.; Goetz, A. W.; Harris, J.; Huang, Z.; Izadi, S.; Izmailov, S. A.; Kasavajhala, K.; Kaymak, M. C.; Kolossváry I.; Kovalenko, A.; Kurtzman, T.; Lee, T. S.; Li, P.; Li, Z.; Lin, C.; Liu, J.; Luchko, T.; Luo, R.; Machado, M.; Manathunga, M.; Merz, K. M.; Miao, Y.; Mikhailovskii, O.; Monard, G.; Nguyen, H.; O'Hearn, K. A.; Onufriev, A.; Pan, F.; Pantano, S.; Rahnamoun, A.; Roe, D. R.; Roitberg, A.; Sagui, C.; Schott-Verdugo, S.; Shajan, A.; Shen, J.; Simmerling, C. L.; Skrynnikov, N. R.; Smith, J.; Swails, J.; Walker, R. C.; Wang, J.; Wang, J.; Wu, X.; Wu, Y.; Xiong, Y.; Xue, Y.; York, D. M.; Zhao, C.; Zhu, Q.; Kollman, P. A. Amber 2025, *University of California, San Francisco*, **2025**.

(15) Ryckaert, J. P.; Ciccotti, G.; Berendsen, H. J. C. Numerical Integration of the Cartesian Equations of Motion of a System with Constraints: Molecular Dynamics of n-Alkanes. *J. Comput. Phys.* **1977**, *23*, 327–341.

(16) Essmann, U.; Perera, L.; Berkowitz, M. L.; Darden, T.; Lee, H.; Pedersen, L. G. A Smooth Particle Mesh Ewald Method. *J. Chem. Phys.* **1995**, *103*, 8577–8593.

(17) Loncharich, R. J.; Brooks, B. R.; Pastor, R. W. Langevin Dynamics of Peptides: The Frictional Dependence of Isomerization Rates of N-acetylalanyl-N′-methylamide. *Biopolymers* **1992**, *32*, 523–535.

(18) Berendsen, H. J. C.; Postma, J. P. M.; Van Gunsteren, W. F.; Dinola, A.; Haak, J. R. Molecular Dynamics with Coupling to an External Bath. *J. Chem. Phys.* **1984**, *81*, 3684–3690.

(19) Roe, D. R.; Cheatham, T. E. I. PTRAJ and CPPTRAJ: Software for Processing and Analysis of Molecular Dynamics Trajectory Data. *J. Chem. Theory Comput.* **2013**, *9*, 3084–3095.

(20) Pettersen, E. F.; Goddard, T. D.; Huang, C. C.; Couch, G. S.; Greenblatt, D. M.; Meng, E. C.; Ferrin, T. E. UCSF Chimera–A Visualization System for Exploratory Research and Analysis. *J. Comput. Chem.* **2004**, *25* (13), 1605–1612.

(21) Meng, E. C.; Goddard, T. D.; Pettersen, E. F.; Couch, G. S.; Pearson, Z. J.; Morris, J. H.; Ferrin, T. E. UCSF ChimeraX: Tools for Structure Building and Analysis. *Protein Sci* **2023**, *32*, e4792.

(22) Ma, S. M.; Tang, Y. Biochemical Characterization of the Minimal Polyketide Synthase Domains in the Lovastatin Nonaketide Synthase LovB. *FEBS J.* **2007**, *274*, 2854–2864.
